# Tuned to explore: Increased phasic responses to auditory targets and novelty in children regardless of induced tonic arousal

**DOI:** 10.1101/2025.11.01.684607

**Authors:** Anne Mathieu, Ursula Schöllkopf, Sophie Chen, Andreas Widmann, Nicole Wetzel, Aurélie Bidet-Caulet

**Author notes:** Equally contributing first authors (Anne Mathieu, Ursula Schöllkopf). Equally contributing last authors (Nicole Wetzel, Aurélie Bidet-Caulet).

## Abstract

The ability to focus on relevant information while ignoring distractions is critical during childhood, as it supports learning, social interaction, and adaptation to changing environments. This attentional balance is thought to depend in part on arousal regulation, mediated by the activity of the locus coeruleus norepinephrine (LC-NE) system. Moderate levels of arousal are typically associated with optimal cognitive performance. However, the interaction between arousal and attention remains poorly understood in humans, especially during development. In this study, we investigated whether experimentally modulating tonic arousal, the baseline level of physiological alertness, affects attentional processing in children (N = 44, aged 6–8) and adults (N = 46, aged 18–35). Participants performed an active auditory three-stimulus (standard, novel, target) oddball task, designed to assess selective attention to target tones and distraction by novel sounds. Prior to each task block, tonic arousal was manipulated using music or videos varying in arousing content. Physiological responses were recorded continuously (skin conductance, pupil dilation, heart rate) to index both tonic arousal and transient, phasic changes in arousal triggered by task events. While tonic arousal modulation was successful, as confirmed by skin conductance levels, Bayesian analyses provided evidence for no effect of this modulation on subsequent attentional processing. Importantly, children generally exhibited stronger phasic arousal responses, particularly to task-irrelevant novel sounds, reflecting less mature regulation of attention and arousal. These findings show developmental differences in physiological responses to unexpected environmental stimuli and provide physiological evidence of increased distractibility during childhood.

## 1 Introduction

Attentional control is the ability to regulate voluntary and involuntary attentional processes, enabling individuals to efficiently filter relevant from irrelevant information. This ability is particularly crucial during childhood, as it underpins the development of core skills required for learning, social interactions, and adapting to complex environments. Voluntary attention allows children to deliberately direct their focus toward aspects of their surroundings in order to achieve specific goals. At the same time, involuntary attentional capture, triggered by unexpected or salient stimuli, provides the flexibility needed to react to surprising changes or novel events in the environment.

### 1.1 Attention: Distraction and facilitation effects

A widely used paradigm to study voluntary and involuntary attentional mechanisms is the three-tone oddball task. Participants are presented with a stream of frequent standard sounds that is interrupted by the infrequent presentation of either (1) target sounds that require a behavioral (voluntary) response or (2) novel sounds that do not require a response and are expected to capture involuntary attention. Involuntary allocation of attention to novel stimuli can be behaviorally measured by a cost in performance (longer reaction times and reduced accuracy), named the **distraction effect** (Bidet-Caulet et al., 2015; Escera et al., 2000; Schröger & Wolff, 1998; Wetzel & Schröger, 2014). Theories suggest that the lengthening of reaction times after the presentation of a novel sound stems from the limited capacity of the attentional system, in which the resources required by the orienting response and the evaluation of the distracting stimulus are then unavailable to perform the task at-hand (Näätänen, 1992). Interestingly, unexpected stimuli can also result in a behavioral benefit, and several studies have reported improved performance when novel sounds are presented (SanMiguel et al., 2010; Wetzel et al., 2012), particularly in the context of highly arousing emotional stimuli (Bonmassar et al., 2023; Max et al., 2015). This **facilitation effect** has been attributed to an increase in phasic arousal triggered by the distracting sound (Grandjean et al., 2025; Masson & Bidet-Caulet, 2019).

### 1.2 Arousal: Physiological level

Arousal can be defined as a level of physiological reactivity that modulates cognitive performance and behavior. Physiological manifestations of arousal are numerous and mediated by the autonomic nervous system (ANS), which include two complementary branches: the sympathetic (SNS) and the parasympathetic (PNS) nervous systems (Karemaker, 2017). Activation of the SNS prepares the body for action by increasing heart rate, dilating pupils, and inhibiting digestion (“fight or flight”), whereas the PNS promotes relaxation by lowering heart rate and stimulating digestion (“rest and digest”). This dynamic balance between the SNS and PNS allows the ANS to adjust the arousal level according to task demands or environmental factors (Wehrwein et al., 2016). Obtaining objective measure of the arousal level remains a challenge. In humans, skin conductance (SC), heart rate (HR) and pupil dilation have been widely used. Indeed, SC reflects the activation of the SNS and is closely linked to emotional arousal (Shehu et al., 2023). The HR is tightly regulated by the dynamic balance between SNS and PNS activity. An increase in HR typically signals SNS activation, whereas a decrease in HR reflects PNS influence (e.g. Wass et al., 2015). Pupil dilation also serves as an indicator of autonomic activity. Tonic as well as phasic increases in pupil size can either be mediated by an increase in SNS activity leading to heightened activity of the dilator muscle or by an inhibition of the PNS leading to a decrease in constriction of the sphincter muscle (McDougal & Gamlin, 2015). In the oddball paradigm, rare or novel sounds elicit a phasic pupil dilation response (PDR; e.g. Wetzel et al., 2016). It has been suggested that the biphasic shape of the PDR is due to the overlap of both the SNS and PNS contribution (Steinhauer & Hakerem, 1992), with the early component reflecting parasympathetic inhibition and the late component reflecting sympathetic activation (Wetzel et al., 2016; Widmann et al., 2018). In an auditory oddball task, only the late component appears to be modulated by the emotional arousing content of the stimulus (Widmann et al., 2018).

### 1.3 Arousal: Brain level

At the brain level, arousal can be described as a general state of cortical excitability, modulated by norepinephrine release from the locus coeruleus (LC; Aston-Jones & Cohen, 2005; Nieuwenhuis et al., 2011). Most findings on the activity of the locus coeruleus – norepinephrine (LC-NE) system come from animal studies (for review see: Aston-Jones & Cohen, 2005). Neurons in the LC exhibit two distinct firing patterns: tonic (sustained) activity, which defines the baseline arousal level and changes relatively slowly, and phasic (transient) activity, which corresponds to bursts of high-frequency firing (10–15 spikes per second) in response to a salient stimulus (Aston-Jones & Cohen, 2005; Vazey et al., 2018). In Aston-Jones and Cohen’s framework, the relation between tonic and phasic LC activity follows an inverted U-shaped curve known as the Yerkes-Dodson relationship (Yerkes & Dodson, 1908): phasic bursts of activity are predominant in a moderate tonic state. Interestingly, behavioral performances in animals follow a similar relationship with LC activity: tonic firing of LC neurons is relatively low when the organism is at rest, medium when engaged in a focused task requiring filtering of irrelevant information, and high when exploring the environment requires staying alert to potentially unexpected occurring events. Thus, a moderate tonic state corresponds to a state of optimal focus and low distractibility (Aston-Jones et al., 1991, 1999; Usher et al., 1999). This observation suggests a strong link between arousal and attentional processes, as advanced by theoretical and physiological models of attention (Broadbent, 1971; Corbetta et al., 2008; Corbetta & Shulman, 2002; Kahneman, 1973; Mesulam, 1981; Posner & Petersen, 1990).

The Adaptative Gain Theory (AGT; Aston-Jones & Cohen, 2005; Gilzenrat et al., 2010; R. Huang & Clewett, 2024) provides a mechanistic account of the arousal-attention relationship, suggesting that variations in arousal modulate neural gain, a process by which high-priority (i.e. task-relevant) signals are enhanced, while low-priority inputs are suppressed. Neural gain is implemented via phasic LC activity generating local “NE-hotspots” (as described in the GANE model; Mather et al., 2016), where high-priority inputs signaled by glutamate are selectively amplified. Recent work integrates the AGT and the GANE model with Kahneman’s capacity model of attention by identifying the LC-NE system as the mechanism that links arousal to both attentional capacity and selectivity (R. Huang & Clewett, 2024). In this view, tonic LC activity mobilizes resources that increase overall processing capacity, while phasic responses sharpen selectivity by amplifying high-priority signals and suppressing distractors. This integration also clarifies that the inverted-U relationship between arousal and performance applies primarily to difficult tasks, in which excessive tonic activity can constrain phasic signaling and excessively narrow selectivity, whereas performance in easier tasks tends to improve more linearly with arousal (R. Huang & Clewett, 2024).

Importantly, the LC is connected to both ANS branches, directly to the SNS and via the Edinger-Westphal nucleus to the PNS. Therefore, physiological measures of arousal have been used as proxy of the LC-NE activity in humans (for review see Reid et al., 2025). Since skin conductance, pupil dilation and heart rate reflect different aspects of ANS activity the combination of all measures can be advantageous, especially in the context of arousal regulation and its impact on voluntary and involuntary attentional mechanisms: The relatively rapid course of the PDR gives precise insight on phasic arousal while tonic SC and HR track (slower) shifts of arousal levels. Due to their non-invasive and sensitive nature, these methods are attractive tools for studying developmental populations.

### 1.4 Developmental perspective

There have been few studies directly comparing the arousal measures in children and adults. Baseline pupil diameter follows a non-linear trajectory across age: it rises in early childhood, peaks in adolescence, and declines thereafter, likely reflecting reduced sympathetic and increased parasympathetic influence in adulthood (Bufo et al., 2022; J. Huang et al., 2024; MacLachlan & Howland, 2002; Tekin et al., 2018; Winston et al., 2020). Regarding skin conductance, children exhibit more frequent spontaneous peaks and a globally larger skin conductance amplitude compared to adults, indicating higher sympathetic activity at rest (Bufo et al., 2022). This suggests that the SNS plays a more active role in childhood which gradually diminishes with maturation, potentially contributing to improved arousal regulation over time. Studies on heart rate maturation similarly indicate higher autonomic activity in children, with faster heart rates compared to adults (Bufo et al., 2022; Finley & Nugent, 1995; Harteveld et al., 2021).

The changes in arousal measures during childhood are closely linked to the progressive development of cognitive functions. Attentional abilities gradually develop throughout childhood, following distinct trajectories for different components of attention. While some aspects of voluntary attention, such as voluntary orienting, are already stable in young infants (Hoyer et al., 2021; Rueda et al., 2004), other aspects like sustained attention appear to develop until early adolescence (Hoyer et al., 2021). Regarding involuntary attentional capture, several studies indicate that children are more distractible than adults, as reflected by larger behavioral costs following the presentation of task-irrelevant sounds. The auditory oddball paradigm is a widely used method for studying the development of deviance detection and distraction control across childhood (for review: Ridderinkhof & van der Stelt, 2000). Behavioral studies consistently show increased reaction times and reduced accuracy to targets following distracting stimuli compared to standard stimuli in younger children compared to older children and adults (Volkmer et al., 2022; Wetzel et al., 2016, 2021). Younger children exhibit a stronger orienting response to unexpected sounds, which leads to greater task disruption. However, with increasing age, they develop better control over these involuntary shifts of attention, resulting in a more stable performance (Volkmer et al., 2022; Wetzel et al., 2021). While detecting novel and unexpected events remains crucial for adapting to a changing environment, the ability to suppress distraction allows older children to better maintain focus on goal-directed tasks, marking a key milestone in cognitive development.

Despite extensive research on attentional development in childhood, relatively few studies have investigated the relationship between attention and arousal. Recent theories suggest that dysregulation of the arousal system may play a role in Attention Deficit Hyperactivity Disorder (ADHD; del Campo et al., 2011). Therefore, some investigations have explored how arousal influences attentional performance in children with ADHD (Bellato et al., 2023; Egeland et al., 2023; Kleberg et al., 2020), indicating that the disorder is often characterized by tonic hypo-arousal, which can sometimes be compensated by heightened phasic responsivity. During typical development, some studies showed correlations between individual tonic arousal level and physiological markers of attention. Wass and collaborators (Wass et al., 2019) used heart rate as an index of tonic arousal during a passive auditory oddball paradigm in 5-7 years old children. Using EEG recording, they looked at the brain responses to relevant and irrelevant stimuli as a function of the arousal level. They showed that children with higher physiological arousal presented larger amplitude P150/P3a responses to deviant tones, suggesting a greater tendency to involuntary attention. One study showed that music tempo affects children attentional performance in an executive attention task (Quan et al., 2023), with faster RT observed during the slow-tempo music compared to fast-tempo music and ocean waves sound. However, subjective ratings of the emotional status of participants did not differ across music conditions, leading the authors to conclude that the behavioral effects were not related to arousal or mood. To our knowledge, no study has successfully manipulated tonic arousal to investigate the resulting effects on attentional performance of children. Therefore, the developmental trajectory of attention and arousal interactions in typically developing children remains largely unexplored, particularly in the auditory modality, limiting our understanding of atypical development.

### 1.5 Aim and Hypothesis

On the basis of current knowledge, we can posit that the still maturing arousal system can differently impact children’s and adults’ reaction to rare relevant and irrelevant stimuli. In the present study, we aim at exploring voluntary and involuntary attention in children and adults under various tonic arousal conditions. For this purpose, we used a three-stimulus oddball paradigm with behavioral and physiological recordings (skin conductance, pupil dilation and heart rate) in 6 to 8 years old children and adults. To our knowledge, this is the first study to simultaneously combine these physiological measures with behavioral performance to investigate attention in young children. Before each block of the oddball task, music and videos were presented during an induction period in order to modulate the tonic arousal level.

We expect generally higher tonic measures (skin conductance, pupil diameter, and heart rate) in children compared to adults, with increased activity for high-arousing conditions in both age groups. During the oddball task, we anticipate: (1) at the behavioral level, greater distraction effects in children, reflected by longer reaction times and higher error rates, and (2) at the physiological level, heightened phasic responses (pupil dilation and skin conductance responses) in children to novel and target sounds, reflecting increased reactivity to salient auditory events. These phasic effects are anticipated to follow the Yerkes-Dodson relationship, where moderate tonic arousal may lead to optimal performance and larger phasic responses to task-relevant stimuli. Whether there are age-related differences in how tonic arousal modulations impact attention remains an open question.

## 2 Methods

### 2.1 Participants

108 participants took part in the experiment. Data from 18 participants had to be discarded because of poor data quality mainly caused by excessive movement, or attention questionnaire scores indicative of symptoms consistent with ADHD. In the end, the analysis was performed on data from **44** children (27 female, mean age ± standard deviation (SD): 7.0 ± 0.8 years) and **46** adults (34 female, mean age ± SD: 24.3 ± 4.3 years). For this collaborative work, half of the data (children: N=24; adults: N=26) were recorded in Magdeburg, Germany, and the other half (children: N=20, adults: N=20) in Marseille, France. Children and adults’ groups were paired in gender and laterality distributions, as well as in socio-economical background. All details regarding participant characteristics can be found in **Table 1** and **Sup Table 1**. All participants were free from neurological or psychiatric disorder, did not take any medication that could impact brain functioning, and had normal hearing and normal or corrected-to-normal vision. The study was approved by a national ethical committee (Marseille) or local ethical committee (Magdeburg), and subjects gave written informed consent (both children, their parents and adults) according to the Declaration of Helsinki. To limit inter-individual differences in tonic arousal, only participants drinking no more than two cups of caffeinated drink and one glass of alcohol per day were included.

**Table 1.** Demographic data of participants.

|  |  | children | adults | age effect |
| --- | --- | --- | --- | --- |
| N |  | 44 | 46 | NA |
| AGE |  | 7.0 | 24.2 | NA |
| GENDER (% female) | | 62 | 74 | $BF_{10} = 0.533$ |
| HANDEDNESS (% left-handed) | | 7 | 15 | $BF_{10} = 0.347$ |
| ATTENTION SCORE |  | 10.7 | 22.0 | NA |
| ED. LEVEL in %<br>(adults' own;<br>parents' for children) | BAC/Abitur | 0 | 2 | $BF_{10} = 0.019$ |
|  | Apprenticeship | 4 | 2 |  |
|  | 2+ years of Uni | 96 | 96 |  |

### 2.2 Stimuli, Software and Presentation

#### Stimuli for oddball task

The three-stimulus oddball task (**Fig. 1**) involved the presentation of three different sounds: standard, target and novel sounds. The complex standard and target sounds consisted of sinusoids with a fundamental frequency of 500 Hz for standards and 1000 Hz for targets; including the second and third harmonic attenuated by −3 and −6 dB, respectively. 32 environmental novel sounds were collected from a database of a previous study (Max et al., 2015). Novel sounds had been rated on a 9-point-scale for valence (1 unpleasant - 9 pleasant) and arousal in adults (1 calm - 9 arousing) (Max et al., 2015). The present study only included moderately arousing neutral sounds (mean ± standard error of the mean (SEM); M_arousal_ = 4.94 ± 0.08; M_valence_ = 5.17 ± 0.09), for example water dripping or clinking glasses. Three additional novel sounds (pig, laughter, squeak) were used for the training phase and were not part of the experiment. All sounds had a duration of 200 ms including 5 ms rise- and fall-time and were equalized using root mean square normalization. Sounds were presented at a loudness of 52.6 dB SPL (measured with PAA3 PHONIC Handheld audio analyzer, Phonic Corporation, Taipei, Taiwan).

**Fig. 1.**
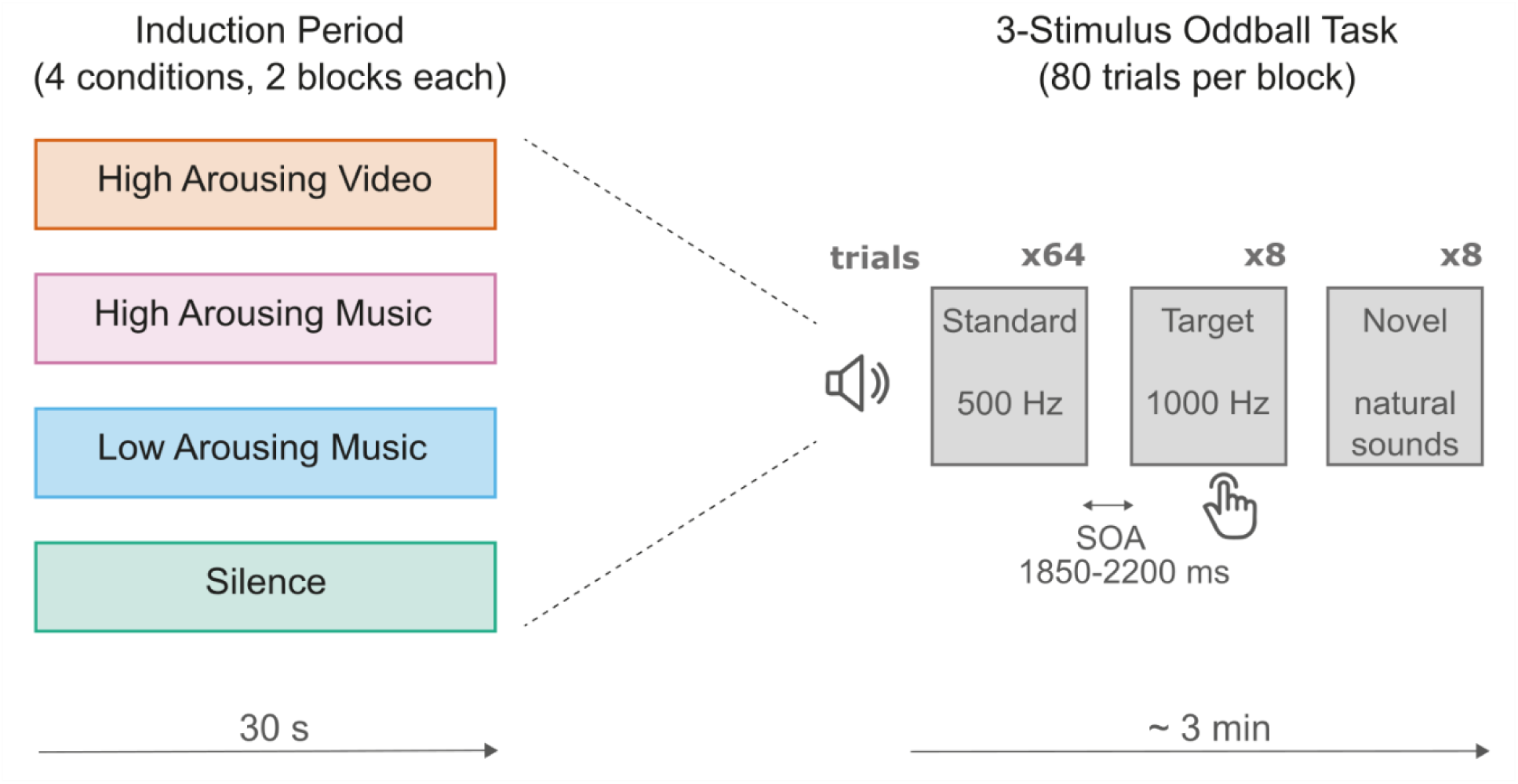
Experimental design. The three-stimulus oddball task involved the presentation of three different sounds: Standard, Target and Novel sounds. Each block of the task (8 in total) was preceded by an induction period during which four types of stimuli were presented: High Arousing Video, High Arousing Music, Low Arousing Music, and Silence. SOA: stimulus-onset asynchrony.

#### Stimuli for tonic arousal modulation

To modulate tonic arousal, musical excerpts rated as low or high arousing were presented at the beginning of each block. The 30 s music excerpts (50 ms rise-time and 50 ms fall-time) were extracted from pieces of classical music categorized and rated in Trost et al. (2012) as function of their arousing and emotional properties. Only excerpts categorized as emotionally positive were chosen (valence superior to 6, from a 10-point scale with 0 = low pleasantness and 9 = high pleasantness). For the present study, music excerpts were divided into two categories: low arousing (mean ± SEM; lowA; M_arousal_ = 2.75 ± 0.15) and high arousing (highA; M_arousal_= 6.32 ± 0.18). Two lowA excerpts were selected from the peacefulness and tenderness categories with an arousal content lower than 3 (10-point scale with 0 = very calming and 9 = very arousing). Four highA excerpts were selected from the wonder, joy and power categories with an arousal content higher than 6. Excerpts were re-sampled at 44100 Hz on 16 bits, edited to last 30 s (50 ms rise-time and 50 ms fall-time), converted from stereo to mono and equalized using root mean square normalization. To amplify the contrast between music conditions, the low and high arousing excerpts were binaurally presented in headphones at 50 dB SPL and 60 dB SPL, respectively. To increase the arousal content, two of the highA musical excerpts were combined with two different cartoon videos that have been selected from a collection of 30 videos that have been previously rated on a 9-point scale for valence (1 happy - 9 neutral) and arousal (1 calm - 9 exciting) by 26 adult participants. Videos were selected for the rating if they could be attributed to the category non-verbal humor and were appropriate for children. The two videos with the highest arousal ratings were selected (mean ± SEM; squirrel from ice-age: M_arousal_ = 5.62 ± 0.32 and cracké bird: M_arousal_ = 5.04 ± 0.30) and adjusted to a mean luminance of 30 +-2 cd/m^2^.

#### Software and presentation

In both labs, the experiment was conducted in an acoustically attenuated shielded cabin. Illuminance of the cabin was held constant at a level of 560 lx in Marseille, 575 lx in Magdeburg (measured with MAVOLUX 5032B USB, GOSSEN Foto- and Lichtmesstechnik GmbH, Nürnberg, Germany). This marginal difference between the two labs likely reflects differences in room dimensions, which can affect how ambient light disperses despite using similar lighting equipment. Participants sat in front of the screen with a viewing distance of 55 cm. The head position was fixed by using a chinrest (SR research head support, Ottawa, Ontario, Canada).

Auditory stimuli were presented via headphones (AKG K142 HD). Visual stimuli in Magdeburg were presented on a 23.6 inch VIEWPixx EEG display (VPixx Technologies Inc.) with a resolution of 1920×1080 (23.6-in. diagonal display size) and a refresh rate of 120 Hz. Visual stimuli in Marseille were presented on a 24.5 inch ACER Predator XN253Q LCD monitor with a resolution of 1920×1080 and a refresh rate of 60Hz. To ensure equal visual stimulus presentation in both labs, a luminance profile of both monitors was measured, and stimuli were adjusted in luminance accordingly. The experimental stimulation was presented via Psychtoolbox (Version 3.0.15, Brainard, 1997). Responses were recorded using a MilliKey response box (MilliKey MH-5 1000 Hz USB).

### 2.3 Task and Procedure

To keep effects of varying instructions in both labs to a minimum, the task was introduced using an introduction video. Participants were instructed to press a button with the index finger of their dominant hand as fast as possible in response to the target sound and were asked to not respond to any other sounds. Target and novel sounds were presented with a 10 % probability. Participants performed a total of eight blocks (80 trials each). In each block, 64 standard sounds, 8 novel sounds and 8 target sounds were presented, resulting in a total of 512 standard sounds, 64 novel sounds and 64 target sounds across the entire task. Sounds were presented in pseudo-randomized order with a randomized sound onset asynchrony (SOA, 1850-2200 ms, 50 ms steps). Each novel and target sound were preceded by at least two standard sounds.

At the beginning of each block, tonic arousal was modulated according to four conditions: (1) low arousing music (LAM), (2) high arousing music (HAM), (3) high arousing music and high arousing video (HAV) (4) silent control condition (S). Two blocks were presented sequentially for each induction condition. The condition order was balanced across participants using latin squares. The task was embedded in a child-friendly story and combined with a game-like visual feedback at the end of each block. Illustrations were displayed at the center of a screen with a size of 1280 x 720 px, 355 x 200 mm (35.8° x 20.6° visual angle from a viewing distance of 550 mm). Mean luminance of the fixation picture was 24 cd/m^2^ on a black background screen with a mean luminance of 0.3 cd/m^2^.

Each experimental block started with a five-point eye-tracker calibration and validation procedure. A baseline period, during which no sound was played, was applied before and after the arousal induction. Before the start of the oddball task, participants were given a short, prerecorded block instruction reminding them to press as fast as possible and to keep their eyes on the screen. The visual feedback on correct button presses was presented after every block. In total, each block lasted around 4 min.

At the beginning of the experiment, participants performed a short training block (21 trials, 3 novels, 3 targets) to get familiarized with the sounds. Before the experimental session, participants filled out a short questionnaire about their socio-demographic characteristics. Adult participants completed the adults ADHD Self-Report Scale (ASRS, Kessler et al., 2005) and parents of children participants completed the ADHD Rating Scale IV (ADHD-RS, DuPaul et al., 1998) assessing attention difficulties. The mean scores (adults: 22.0 ± 8.1; children: 10.7 ± 6.4) confirmed our population did not present attentional difficulties. In addition, a sleep questionnaire was fulfilled at the beginning of each experimental session to assess the sleep quality on the night before experimental session. All participants stated that they had satisfactory slept the night before. After the experimental session, participants were asked to rate the music and video excerpts that have been used in the experiment, on a 9-point scale for valence (1 happy - 9 sad) and arousal (1 calm - 9 exciting).

### 2.4 Data recording

The pupil diameter of both eyes was recorded during each entire block (arousal induction + task) with an infrared EyeLink Portable Duo (SR Research, Ottawa, Ontario, Canada) remote eye tracker at a sampling rate of 500 Hz. Participants were instructed to blink naturally during the task but to keep their eyes on the screen during the task.

For skin conductance (SC), electrodes were placed on the medial phalanx of the index and middle fingers of the non-dominant hand. For electrocardiogram (ECG), electrodes were placed on the left shoulder and stomach. In Magdeburg, SC and ECG responses were measured with the GSR module and the BIP2AUX adapter from an ActiChamp amplifier at a sampling rate of 500 Hz, respectively (Brain Products GmbH, Gilching, Germany). In Marseille, Flat-Type active electrodes (ActiveTwo system, BioSemi, Netherlands) were used. Data was then amplified (−3 dB at ∼204 Hz low-pass, DC coupled) and digitized (1024 Hz).

### 2.5 Data analysis

The first two standard trials per block are important to establish a standard sound representation for the following stimuli and were thus excluded (Bendixen et al., 2007). The two standard trials following a novel or a target sound can be affected by preceding distractor processing (Wetzel, 2015) or the response to the target and were also excluded from all analyses. This also promotes enough time for the pupil to return to baseline after a response or a novel sound. Only trials where participants’ responses were given correctly and in the time-window from 300 ms to 1700 ms after sound onset were used for behavior, skin conductance and pupil analysis.

#### Behavioral measures

In target trials, no button press in the set response time-window (300 ms – 1700 ms) was considered a Miss. In standard and novel trials, a button press between the RT lower and upper limit was considered a False alarm (FA), no button press was considered a Correct rejection (CR).

#### Skin conductance

Skin conductance data analysis was performed using the software package for electrophysiological analysis (ELAN Pack) developed at the Lyon Neuroscience Research Center (; Aguera et al., 2011), Ledalab, a free and open-source Matlab-based program used for the analysis of skin conductance data (Benedek & Kaernbach, 2010) and custom MATLAB programs.

Data were low pass filtered offline at 10 Hz. Data recorded in Marseille was resampled at 500 Hz to match the sampling frequency of data recorded in Magdeburg. Continuous phasic and tonic activities were extracted using the Continuous Decomposition Analysis tool from Ledalab. For phasic activity, a baseline subtraction was applied using the 500 ms period before stimulus onset, and amplitudes of the skin conductance responses (*SCR*) to standard, novel and target sounds were computed in the time-window from 1500 to 5000 ms after sound onset. This window corresponds to the typical SCR latency range (Boucsein et al., 2012; Dawson et al., 2000) and was confirmed by inspecting individual data to capture the peak response in both children and adults. For modulations of the tonic activity, a baseline subtraction was applied using the 5000 ms period before induction period onset, and skin conductance level (*SCL*) was computed in 15 s time-windows from 0 to 195 s after induction onset (0-15; 15-30; 45-60; 60-75; 75-90; 90-105; 105-120; 120-135; 135-150; 150-165; 165-180; 180-195).

#### Heart rate

Electrocardiogram data analysis was performed using custom MATLAB programs. ECG data were filtered offline using a 1 Hz high-pass filter and a 100 Hz low-pass filter (MATLAB). R-peaks were automatically detected, and their detection was validated through a visual sanity check to identify any missing, duplicate, or incorrectly placed peaks. Any mis-detected peak was manually corrected using Anywave software (Colombet et al., 2015). Interbeat intervals (IBIs) were then derived from the signal. Mean IBI were calculated in 30 s time-windows from 0 to 195 s after induction onset (0-15; 15-30; 45-60; 60-75; 75-90; 90-105; 105-120; 120-135; 135-150; 150-165; 165-180; 180-195). The standard deviation of the IBI series in each 30 s time-windows were calculated for each block and participant to quantify heart rate variability.

#### Pupillometry

Pupil data analysis was carried out using MATLAB custom programs. The pupil diameter digital counts of the eye-tracker were calibrated using the method introduced by (Steinhauer et al., 2022) and converted to millimeter. Blinks and enclosing saccades were marked by the blink events provided by the eye-tracker. Partial blinks (not reported by the eye-tracker) were identified from the smoothed velocity time series as pupil diameter changes higher than 20 mm/s including a 50 ms pre-blink and a 100 ms post-blink interval using an additional custom function (Merritt et al., 1994). The data from both eyes were averaged using the dynamic offset algorithm suggested by Kret & Sjak-Shie (2019). Blinks and partial blinks (and intervals with signal loss) shorter than 1 s were interpolated using piecewise cubic Hermite polynomial interpolation. Missing data segments longer than 1 s were discarded from the data. Data was then further processed for analyzing (1) tonic pupil dilation or (2) phasic pupil dilation responses following two different processing streams: (1) For the tonic pupil dilation, continuous data was divided into 12 time-windows (15 s, similar to heat rate and SCL analyses) per block and averaged per participant and arousing condition. (2) For the phasic pupil dilation response (PDR) the continuous data was segmented into epochs of 2 s duration around sound onset (including a −200 ms to 0 ms pre-stimulus baseline). PDR epochs were baseline corrected by subtracting the mean amplitude of the baseline period (200 ms) from each epoch (Murphy et al., 2014; Widmann et al., 2018). Individual average PDRs were computed per participant, sound type (standard, target, novel) and arousing condition. A time-window was centered on the early and late peaks of the averaged dilation responses for each participant (adults: early peak 500-800 ms; late peak 1100-1500 ms; children: early peak 600-900ms; late peak 1000-1500 ms). The time-windows (TW) for the early and late peaks were determined separately for each age group, based on the grand-average waveforms collapsed across all conditions, to capture group-specific peak latencies while avoiding bias from condition-specific variability.

### 2.6 Statistical analysis

Bayesian ANOVAs and Bayesian post-hoc t-tests were conducted using JASP (JASP Team, 2024, version 0.95.0.0). For behavioral (response rates, RT, RT variability and ratings) and tonic measures (SCL, pupil dilation, IBI and IBI variability in TW 1 to 12), age (levels: children, adults) was used as between-subjects and induction (levels: silence (S), low arousing music (LAM), high arousing music (HAM), high arousing video (HAV)) as within-subjects factors. For phasic (PDR and SCR) measures, sound type (levels: standard, target, novel) was added as additional within-subjects factor.

Bayes factor (BF_10_) for the best supported model is reported, where larger values suggest more evidence is provided by the data for the model reflecting the alternative hypothesis or H_1_ (lower for model reflecting the null hypothesis H_0_). For interpretation, the scale provided by Lee & Wagenmakers (2014) was followed, where moderate evidence in favor of H_1_ (or H_0_) is considered if BF_10_ is larger than 3 (or lower than 0.33), strong evidence if BF_10_ is larger than 10 (lower than 0.1), or decisive evidence if BF_10_ is larger than 100 (lower than 0.01). BF_10_ between 0.33 and 3 is considered as anecdotal evidence. BFs were estimated using 100,000 Monte-Carlo sampling iterations. Effect size of the priors for the fixed and the random effects were set to scaling factors of r = 0.5 and r = 1, respectively. Additionally, inclusion Bayes factors based on matched models were calculated by comparing the models containing a main or interaction effect to the equivalent models stripped of the effect (BF_Incl_; Mathôt, 2017). Post-hoc Bayesian independent samples t-tests (for between-subjects factor) and Bayesian paired-samples t-tests (for within-subject factors) were used to analyze main effects and interactions, where the null hypothesis corresponded to a standardized effect size δ= 0, while the alternative hypothesis was defined as a Cauchy prior distribution centered around 0 with a scaling factor of r= 0.707.

## 3 Results

### 3.1 Ratings of induction stimuli

Valence and Arousal ratings of induction stimuli were analyzed using Bayesian ANOVA with the factors Induction and Age.

For Valence ratings, the best-supported model included Induction as predictor (BF_10_ = 2.44 × 10^+26^). There was decisive evidence for an effect of Induction (**Fig 2.A**; BF_incl_ = 3.10 × 10^+26^) and anecdotal evidence for no effect of Age (BF_incl_ = 0.99). Post-hoc Bayesian paired-samples t-tests revealed no difference in valence ratings between High Arousing Music and High Arousing Video stimuli (BF_10_ = 0.112). In contrast, Low Arousing Music was rated as less joyful compared to both High Arousing Music (BF_10_ = 1.49 × 10^+15^) and High Arousing Video (BF_10_ = 4.46 × 10^+14^).

**Fig. 2.**
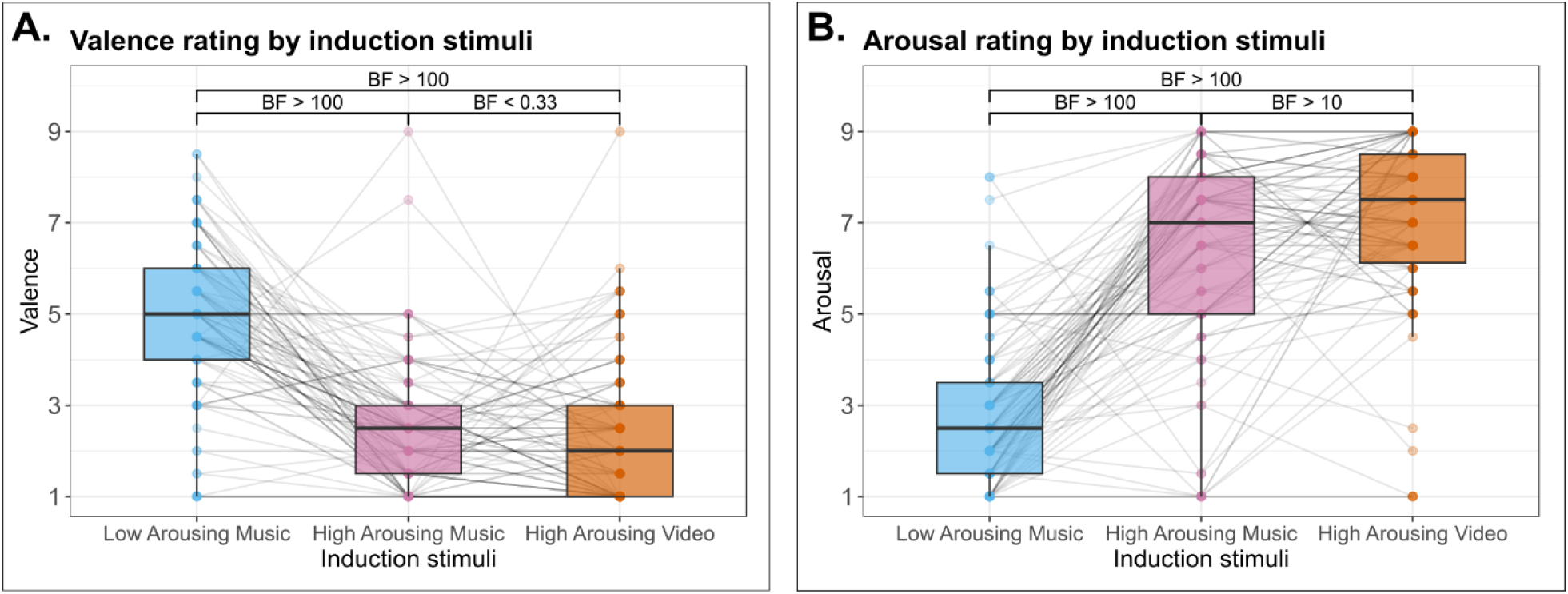
Subjective ratings of induction stimuli. **A**: Valence ratings. **B**: Arousal ratings. Within each boxplot, the horizontal line represents the group median, the lower and upper hinges correspond to the first and third quartiles. The upper whisker extends from the hinge to the largest value no further than 1.5 * IQR from the hinge (IQR = inter-quartile range, or distance between the first and third quartiles). The lower whisker extends from the hinge to the smallest value at most 1.5 * IQR of the hinge. Superimposed to each boxplot, the dots represent individual means. Light grey lines connect the dots belonging to the same participant across conditions. BF: Bayes Factor.

For Arousal ratings, the best-supported model included Induction, Age and Induction x Age as predictors (BF_10_ = 3.14 × 10^+45^). There was strong to decisive evidence for an effect of Induction (**Fig 2.B**; BF_incl_ = 5.59 × 10^+44^) and for an effect of Induction x Age (BF_incl_ = 32.26). There was moderate evidence for no effect of Age (BF_incl_ = 0.179). Post-hoc Bayesian t-tests were conducted to explore the Induction × Age interaction (Sup. Fig 1). Children rated High Arousing Video as more arousing than both Low Arousing Music (BF_10_ = 9.15 × 10^+8^) and High Arousing Music (BF_10_ = 1.49 × 10^+4^), and also rated High Arousing Music as more arousing than Low Arousing Music (BF_10_ = 46.52). In contrast, adults rated High Arousing Video and High Arousing Music as equally arousing (BF₁₀ = 0.153) and rated Low Arousing Music as less arousing than both High Arousing Video (BF_10_ = 2.54 × 10^+21^) and High Arousing Music (BF_10_ = 8.29 × 10^+19^). There was moderate to no evidence for differences between children and adults whatever the induction (BF_10_ <= 3.82). Details of post-hoc comparisons can be found in Sup. Table 2.

In summary, induction stimuli differed in both valence and arousal ratings, with children showing greater sensitivity to arousal differences between high-arousing conditions.

### 3.2 Behavioral data

#### Response rates

The percentage of False Alarms (FA) in standard and novel trials was analyzed using Bayesian ANOVA with the factors Induction and Age. The best-supported model included Age as a predictor (BF_10_ = 1.27 × 10^+3^). There was decisive evidence for an effect of Age (**Fig 3.A**; BF_incl_ = 1.26 × 10^+3^) with children making more FA than adults. In contrast, there was moderate to strong evidence for no effect of Induction (BF_incl_ = 0.038) and for no Age x Induction interaction (BF_incl_ = 0.290).

**Fig. 3.**
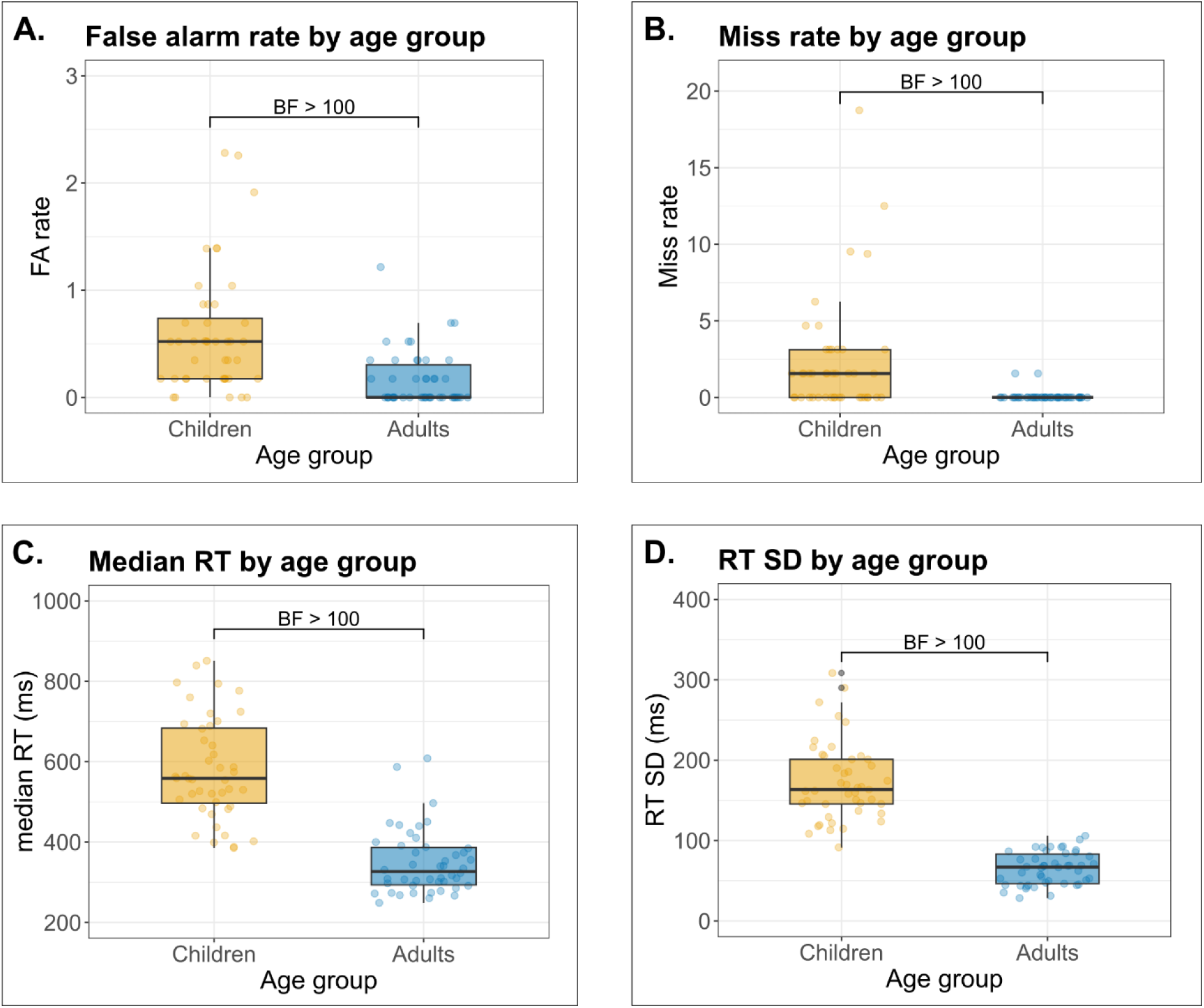
Behavioral results by age group. **A**: False alarm rate. **B**: Miss rate. **C**: Reaction times. **D**: Reaction times variability. Within each boxplot, the horizontal line represents the group median, the lower and upper hinges correspond to the first and third quartiles. The upper whisker extends from the hinge to the largest value no further than 1.5 * IQR from the hinge (IQR = inter-quartile range, or distance between the first and third quartiles). The lower whisker extends from the hinge to the smallest value at most 1.5 * IQR of the hinge. Superimposed to each boxplot, the dots represent individual means. BF: Bayes Factor.

Similarly, the percentage of Misses in target trials was analyzed using Bayesian ANOVA with the factors Induction and Age. The best-supported model again included Age as a predictor (BF_10_ = 190.86). Decisive evidence supported an effect of Age (**Fig 3.B**; BF_incl_ = 191.00), indicating that children missed more targets than adults. In contrast, there was moderate to strong evidence for no effect of Induction (BF_incl_ = 0.070) or of its interaction with Age (BF_incl_ = 0.185).

#### Reaction times

Reaction times to all correctly detected targets were analyzed using Bayesian ANOVA with the factors Induction and Age. The best-supported model included Age as a predictor (BF_10_ = 5.58 × 10^+15^). There was decisive evidence for the effect of Age (**Fig 3.C**; BF_incl_ = 5.58 × 10^+15^) indicating that adults were faster than children to respond to target sounds. There was decisive evidence for no effect of Induction (BF_incl_ = 0.020) and for no Age x Induction interaction (BF_incl_ = 0.035).

#### Reaction times variability

Standard deviation of reactions times to all correctly detected targets was analyzed using Bayesian ANOVA with the factors Induction and Age. The best-supported model included Age as a predictor (BF_10_ = 5.61 × 10^+20^). There was decisive evidence for the effect of Age (**Fig 3.D**; BF_incl_ = 5.62 × 10^+20^) as adults’ reaction times were less variable than those of children. We found strong evidence for no effect of Induction (BF_incl_ = 0.024) and for no Age x Induction interaction (BF_incl_ = 0.031).

To assess whether an induction effect could be limited to the beginning of the block, additional analyses of *Miss*, *FA*, *RT,* and *RT variability* were performed on the first half of trials for each block. Bayesian ANOVAs showed positive to strong evidence for no effect of Induction on behavioral measures (0.037 < BF_incl_ < 0.890).

In summary, compared to adults, children made more false alarms, missed more targets, responded more slowly and showed greater variability in reaction times, with no effect of induction condition.

### 3.3 Physiological indices of tonic arousal

The mean amplitude of physiological indices was analyzed using Bayesian ANOVA with the factors Induction, Age, and Time-window (TW). When evidence for interactions with the TW factor was found, we further performed separate analyses for each of the twelve 15-second TWs corresponding to the induction period (TW-1 and TW-2) and the following oddball task (TW-3 to TW-12).

#### Skin conductance level (SCL)

For the mean amplitude of tonic SCL, we found decisive evidence for interactions between TW and Age (BF_incl_ = 2.46 × 10^+^¹³) and TW and Induction (BF_incl_ = 5.05 × 10^+^⁴). Details of the models for all TWs can be found in Table 2.A. Details of post-hoc comparisons can be found in Sup. Table 3.

**Table 2.**
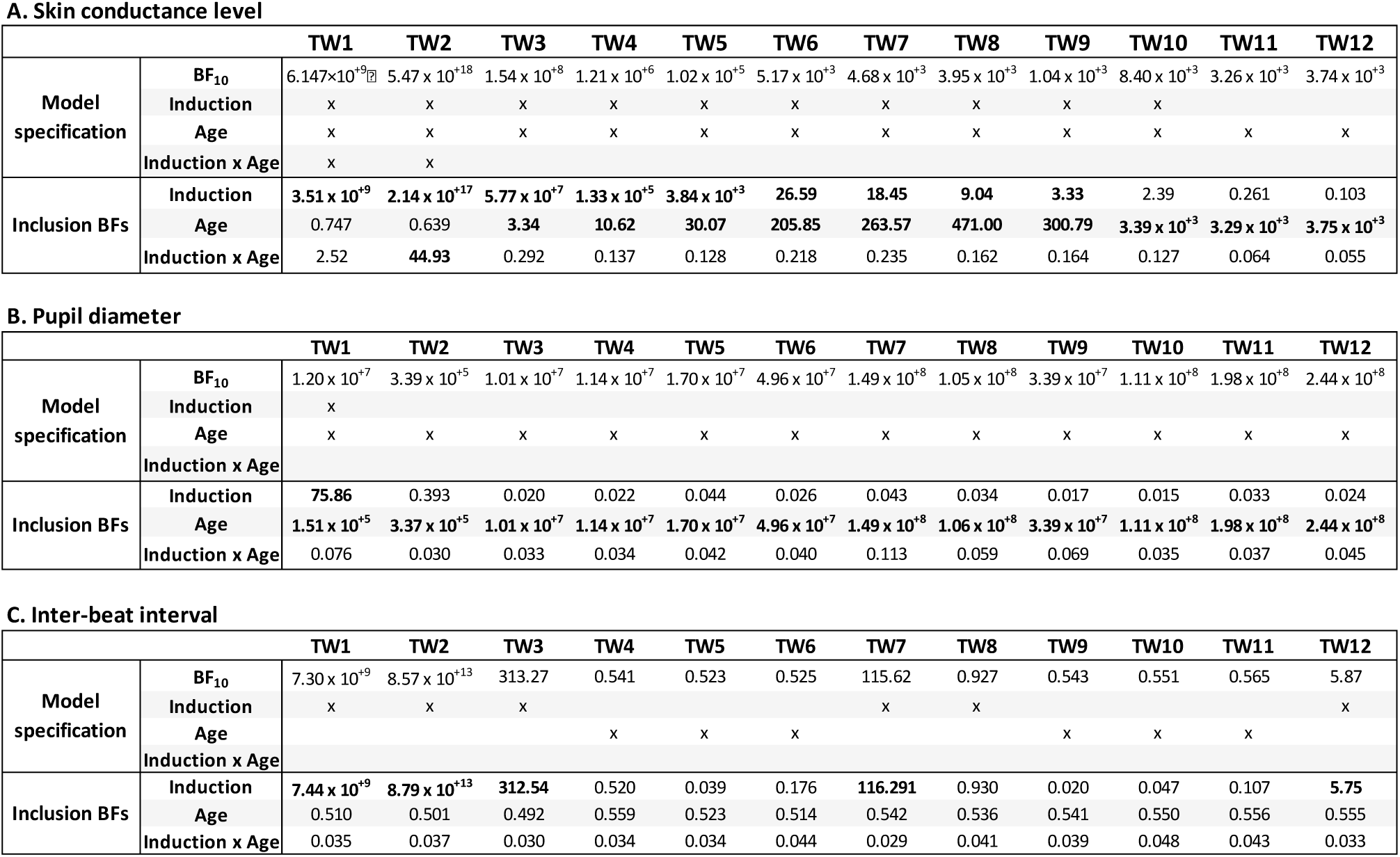
Best model specification and inclusion BFs for Bayesian ANOVAs on tonic measures: skin conductance level (A), pupil diameter (B) and inter-beat interval (C). BF: Bayes Factor.

There was moderate to decisive evidence for the effect of **Induction** in TW-1 to TW-9 (3.33 < BF_incl_ < 2.14 × 10^+17^). Post-hoc Bayesian paired samples t-tests revealed that the High Arousing Video condition elicited higher SCL than all other conditions in TW-1 to TW-5 (all BF_10_ > 7.34), and higher SCL than High Arousing Music in TW-6 to TW-9 (all BF_10_ > 30.57).

There was moderate to decisive evidence for the effect of **Age** in TW-3 to TW-12 (3.34 < BF_incl_ < 3.39 × 10^+3^), with children showing higher SCL than adults.

There was strong evidence for an **Induction × Age interaction** in TW-2 (BF_incl_ = 44.93), with post-hoc tests showing that children had higher SCL than adults in the High Arousing Video condition only (BF_10_ = 6.79; all other BF_10_ < 0.32, see Sup. Fig. 2).

In summary, high-arousing videos increased SCL during the induction period and the beginning of the task (**Fig 4.A**). Additionally, children exhibit larger SCL than adults throughout most of the oddball task (**Fig 4.B**), with a transient increase specifically in response to the high-arousing video during the second half of the induction period.

**Fig. 4.**
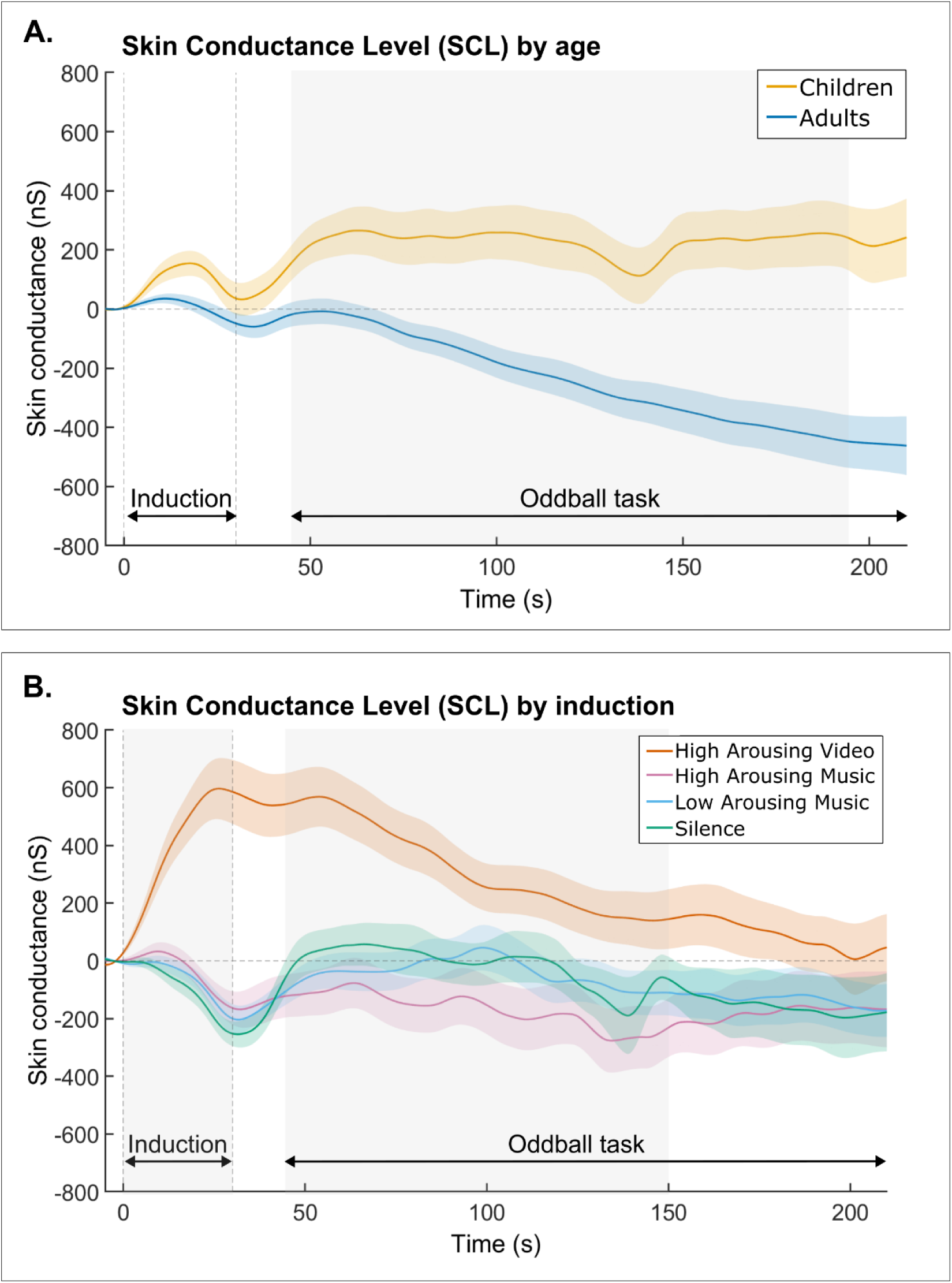
Skin conductance level (SCL) time courses. **A**: SCL by age group. **B**: SCL by induction condition. Shadowed areas represent standard errors of the mean. Grey areas correspond to time-windows with evidence for Age or Induction effect.

#### Pupil size

For the mean amplitude of the pupil size, we found decisive evidence for a TW x Age interaction (BF_incl_ = 1.77 × 10^+14^). Details of the models for all TWs can be found in Table 2.B, and for post-hoc comparisons in Sup. Table 4.

There was decisive evidence for the effect of **Induction** in TW-1 (BF_incl_ = 7.44 × 10^+9^). Post-hoc Bayesian paired samples t-tests revealed that high-arousing conditions (HAV and HAM) elicited larger pupil diameter than silence (BF_10_ > 8.32).

There was decisive evidence for the effect of **Age** in all TWs (1.51 × 10^+5^ < BF_incl_ < 2.44 × 10^+8^), indicating that children presented larger pupil diameter compared to adults.

In summary, children displayed larger pupil size compared to adults throughout both the induction period and the oddball task (Sup. Fig 3). By contrast, the induction condition appeared to have effect on pupil size only during the first 15 seconds.

#### Heart rate

For the mean inter-beat interval (IBI), we found decisive evidence for a TW x Induction interaction (BF_incl_ = 1.05 × 10^+113^). Details of the models for all TWs can be found in Table 2.C, and for post-hoc comparisons in Sup. Table 5.

There was moderate to decisive evidence for the effect of **Induction** in TW-1 to TW-3, in TW-7 and TW-12 (5.75 < BF_incl_ < 8.79 × 10^+13^). Post-hoc Bayesian paired samples t-tests revealed that, in TW-1 to TW-3, the mean IBI in High Arousing Video condition was longer than in all other conditions (all BF_10_ > 11.69), meaning slower heart rate in the High Arousing Video condition. In TW-7 and TW-12, the mean IBI was shorter in high-arousing conditions (HAV; HAM) condition compared to Silence (all BF_10_ > 3.38).

In TW-4 to TW-6 and in TW-8 to TW-11, no model provided stronger support than the null model (including only subject as a random factor, 0.523 < BF_10_ < 0.927).

In summary, the induction effects on mean IBI appear inconsistent, with arousing stimuli either resulting in longer IBI during the induction period or shorter IBI during a brief portion of the oddball task (Sup. Fig 4). Furthermore, age had no impact on the mean IBI.

#### Heart rate variability

The standard deviation of IBI was analyzed using Bayesian ANOVAs with the factors Induction and Age for each of the twelve 15-seconds TW. For TW-3 to TW-7 and TW-9 to TW-12, the best-fitting models included Age (1.13 < BF_10_ < 2.24). However, evidence for the effect of Age was only anecdotal (1.12 < BF_incl_ < 2.24), with children showing greater variability than adults. For TW-1, TW-2 and TW-8, no model provided stronger support than the null model (including only subject as a random factor, 0.644 < BF_10_ < 0.975). For all TWs, there was anecdotal to strong evidence against an effect of Induction on IBI variability (0.016 < BFincl < 0.649).

In summary, IBI variability shows only weak evidence for age-related differences, with no evidence for an influence of induction.

### 3.4 Phasic responses during the oddball tasks

#### Skin conductance responses (SCR)

The mean SCR amplitude (1500 to 5000 ms) in response to sound was analyzed using Bayesian ANOVA with the factors Induction, Age and SoundType. The best-supported model included Age, SoundType and the Age x SoundType interaction (BF_10_ = 2.49 × 10^+18^). There was decisive evidence for the SoundType effect (**Fig 5.B**; BF_incl_ = 3.57 × 10^+12^), the Age effect (**Fig 5.C**; BF_incl_ = 456.30) and the Age x SoundType interaction (**Fig 5.A and 5.D**; BF_incl_ = 1.17 × 10^+3^). Main effects indicated that SCR was largest for target sounds, intermediate for novel sounds, and smallest for standard sounds (all BF_10_ > 1.05 × 10^+9^), and was overall larger in children than in adults. There was also decisive evidence against an Induction effect (BF_incl_ = 0.004).

**Fig. 5.**
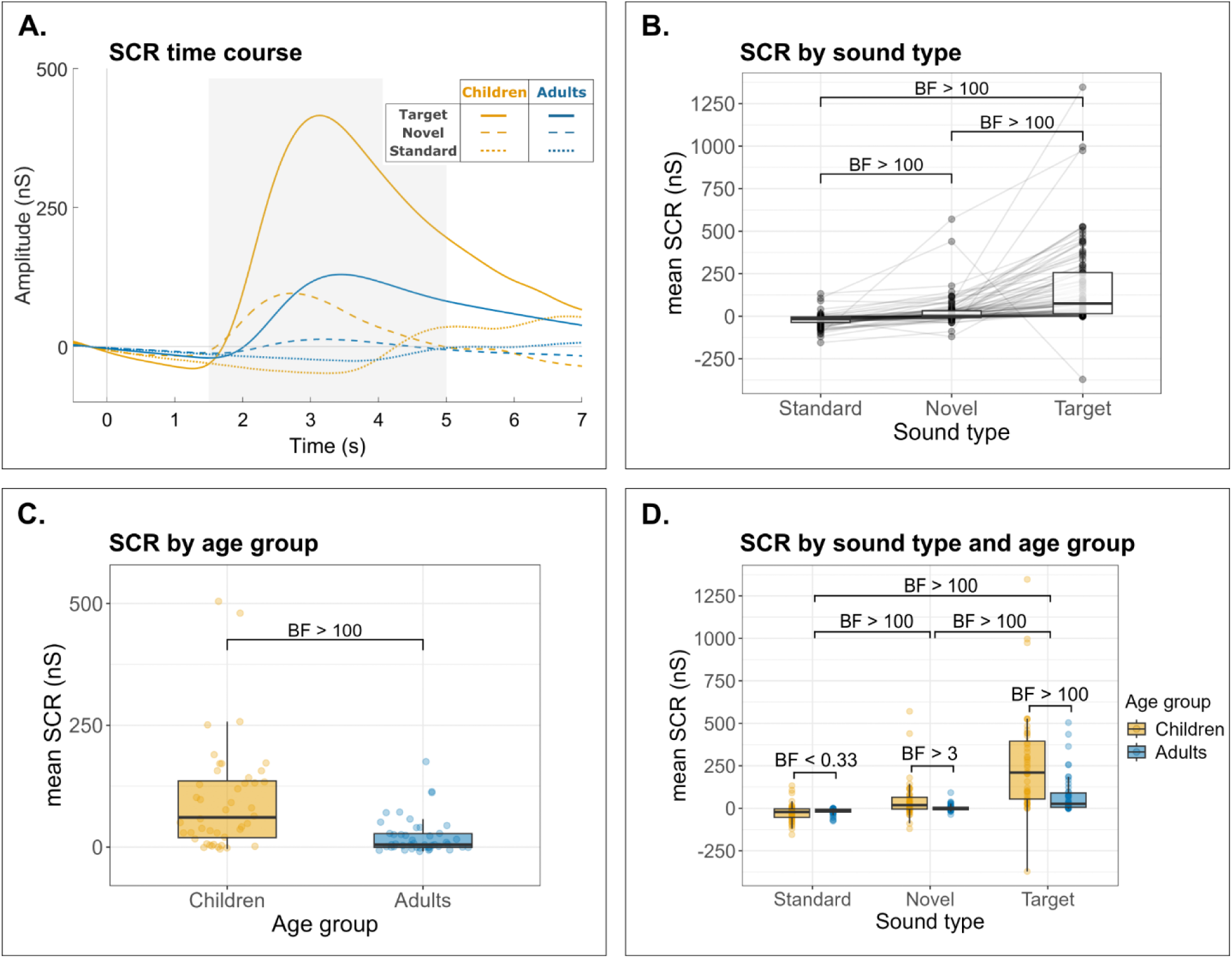
Skin Conductance Response (SCR). **A**: SCR time course by sound type and age group. **B**: mean SCR by sound type. **C**: mean SCR by age group. **D**: mean SCR by sound type and age group. Boxplots in panels **B, C** and **D** were computed using the mean amplitude in the 1.5 – 5 s time-window after sound onset (grey area on the time course). Within each boxplot, the horizontal line represents the group median, the lower and upper hinges correspond to the first and third quartiles. The upper whisker extends from the hinge to the largest value no further than 1.5 * IQR from the hinge (IQR = inter-quartile range, or distance between the first and third quartiles). The lower whisker extends from the hinge to the smallest value at most 1.5 * IQR of the hinge. Superimposed to each boxplot, the dots represent individual means. Light grey lines connect the dots belonging to the same participant across conditions. BF: Bayes Factor.

Post-hoc Bayesian independent samples t-test were used to investigate the Age x SoundType interaction (Sup. Table 6). There was moderate to decisive evidence for an Age effect on the SCR to novel (BF_10_ = 4.20) and target sounds (BF_10_ = 148.83): children presenting larger responses than adults to rare stimuli. In contrast, there was moderate evidence for no difference between children’s and adults’ SCR to standard sounds (BF_10_ = 0.289).

In summary, children showed larger SCR amplitudes than adults for rare salient sounds (novels and targets) but not for standard sounds, while induction had no effect.

To assess whether an induction effect could be limited to the beginning of the block, we analyzed the mean SCR amplitude (1500 to 5000 ms) in response to sounds presented during the first half of the trials. A Bayesian ANOVA was conducted with the factors Induction, Age, and SoundType. The results were similar to those observed for the entire block: the best supported model was SoundType + Age + SoundType x Age (BF_10_ = 6.41 × 10^+20^), with decisive evidence against an Induction effect (BF_incl_ = 0.016). Post-hoc comparisons can be found in Sup. Table 7.

#### Pupil Dilation responses (PDR)

The mean amplitude during the early (adults: 500-800ms, children: 600-900ms) and late (adults: 1100-1500ms, children: 1000-1500ms) peaks of the PDR to sound was analyzed using Bayesian ANOVA with the factors Induction, Age and Sound Type.

For the early peak, the best supported model included Age, SoundType, and the Age x SoundType interaction (BF_10_ = 1.15 × 10^+49^). There was decisive evidence for the SoundType effect (**Fig 6.B**; BF_incl_ = 1.98 × 10^+48^), indicating that the early PDR peak was largest for target sounds, intermediate for novel sounds, and smallest for standard sounds (all BF_10_ > 211.05). There was also decisive evidence for the Age x SoundType interaction (**Fig 6.A and 6.C**; BF_incl_ = 29.81). However, there was moderate evidence against an Age effect (BF_incl_ = 0.197) and decisive evidence against an Induction effect (BF_incl_ = 0.004).

**Fig. 6.**
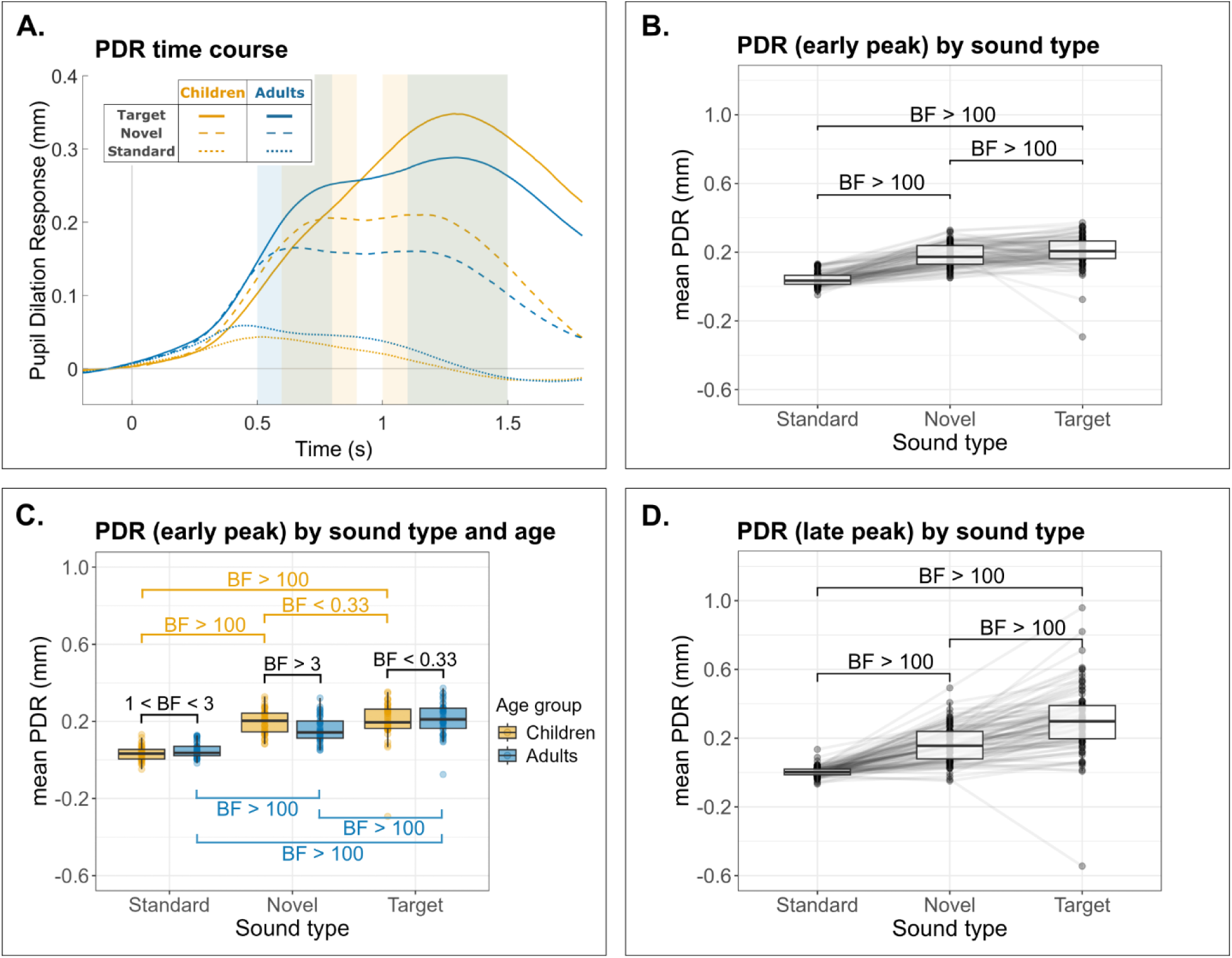
Pupil Dilation Response (PDR). **A**: PDR time course by sound type and age group. Light yellow areas correspond to the time-windows for early (600-900 ms) and late (1000-1500 ms) peaks in children. Light blue areas correspond to the time-windows for early (500-800 ms) and late (1100-1500 ms) peaks in adults. **B**: mean PDR in early peak by sound type. **C**: mean PDR in early peak by sound type and age group. **D**: mean PDR in late peak by sound type. Boxplots were computed using the mean amplitude in the early (**B** and **C**) or late (**D**) peaks time-window. Within each boxplot, the horizontal line represents the group median, the lower and upper hinges correspond to the first and third quartiles. The upper whisker extends from the hinge to the largest value no further than 1.5 * IQR from the hinge (IQR = inter-quartile range, or distance between the first and third quartiles). The lower whisker extends from the hinge to the smallest value at most 1.5 * IQR of the hinge. Superimposed to each boxplot, the dots represent individual means. Light grey lines connect the dots belonging to the same participant across conditions. BF: Bayes Factor.

Post-hoc Bayesian paired-samples t-tests were used to investigate further the Age x SoundType interaction (Sup. Table 6). Contrary to adults presenting increasing PDR amplitude from standard to novel to target sounds (all BF_10_ > 6.97 × 10^4^); in children, there was strong evidence for no difference in amplitude between the PDR to novel and target sounds (BF_10_ = 0.170). Post-hoc independent samples Bayesian t-tests provided moderate evidence for the early peak PDR amplitude in children being larger compared to adults in response to novel sounds (BF_10_ = 6.46), but similar to adults in response to targets sounds (BF_10_ = 0.252).

For the late peak, the best-supported model included SoundType (BF_10_ = 6.21 × 10^+42^). There was decisive evidence for the effect of SoundType (**Fig 6.D**; BF_incl_ = 6.30 × 10^+42^), indicating that the late PDR peak was largest for target sounds, intermediate for novel sounds, and smallest for standard sounds (all BF_10_ > 5.61 × 10^35^). There was anecdotal evidence against an effect of Age (BF_incl_ = 0.710) and against an Age x SoundType interaction (BF_incl_ =0.547), and strong evidence against an effect of Induction (BF_incl_ = 0.010). Details of post-hoc comparisons can be found in Sup. Table 6.

In summary, children displayed a larger PDR early peak than adults in response to novel sounds, reaching an amplitude similar to their response to targets. Induction conditions had no effect on any of the PDR peaks.

To assess whether an induction effect could be limited to the beginning of the block, we analyzed the mean PDR amplitude (early and late peaks) in response to sounds presented during the first half of the trials. Bayesian ANOVAs were conducted with the factors Induction, Age, and SoundType. The results were similar to those observed for the entire block: the best supported model was SoundType + Age + SoundType x Age (BF_10_ = 1.83 × 10^+53^) for the early peak, and SoundType (BF_10_ = 2.33 × 10^+43^) for the late peak, with decisive evidence against an Induction effect (BF_incl_ < 0.019). Post-hoc comparisons can be found in Sup. Table 7.

## 4 Discussion

We aimed at investigating how modulation of tonic arousal affects voluntary and involuntary attention in 6–8 year-old children and adults. Specifically, we explored the influence of musical and video excerpts on behavioral and physiological phasic responses to target and novel sounds within an auditory three-stimulus oddball paradigm.

Skin conductance (SC) and pupil diameter (PD) measures confirmed successful upregulation of tonic arousal. However, Bayesian analyses showed evidence for no effect of induction on phasic responses or behavioral performance in either age group. Crucially, we found strong developmental effects, irrespective of tonic arousal modulation. Children, compared to adults, showed increased phasic responses, with effects depending on the sound type: skin conductance responses (SCRs) were higher for both task-relevant (target) and irrelevant (novel) sounds, while the early peak of the pupil dilation response (PDR) was larger only in response to task-irrelevant sounds. In the following, we will discuss (1) how tonic arousal is challenging to modulate and how it influences performance and phasic physiological responses during the attention task and (2) developmental changes in autonomic responses, focusing on how they reflect voluntary and involuntary attentional mechanisms.

### Arousal induction effects

#### Presentation of audiovisual stimuli modulates physiological measures of tonic arousal

We first assessed whether our induction procedure effectively impacted tonic arousal measures. The large increase in skin conductance level (SCL) during the combined presentation of high-arousing music and video (HAV) indicates successful modulation of tonic arousal in both age groups. In contrast, high-arousing music presented alone failed to induce major changes. This is in line with research on multisensory integration suggesting that the simultaneous presentation of both visual and auditory congruent stimuli can enhance perceptual processing including emotional experience (for review see Opoku-Baah et al., 2021).

Interestingly, the HAV-induction effect was longer lasting for SCL compared to the tonic pupil diameter (PD). Only a transient induction effect during the first 15 sec was found on the PD. This transient effect on tonic PD was also present in the high-arousing music condition. The difference in sensitivity towards tonic arousal modulation may be attributed to the distinct autonomic pathways regulating SCL and PD. SC reflects changes in sweat gland activity and can last for several seconds. It is primarily regulated by the sympathetic nervous system (SNS), which is closely associated with emotional arousal (Boucsein et al., 2012). PD on the other hand is regulated by both sympathetic and parasympathetic inputs and is more closely tied to rapid changes in arousal (Einhäuser, 2017; Murphy et al., 2014; Widmann et al., 2018).

#### Increasing arousal does not affect behavioral or physiological responses during the task

Our findings showed no effect of tonic arousal modulation on responses to standard, target, or novel sounds, at both behavioral and physiological levels, in adults and children.

This result contrasts with comparable studies by Mather et al. (2020) and Meissner et al. (2024), who investigated how reducing tonic LC-NE activity, via isometric handgrip (Mather et al., 2020) or pupil-based biofeedback (Meissner et al., 2024), influenced subsequent pupil dynamics during an auditory oddball task. Both studies observed that a *decrease* in tonic pupil size was associated with enhanced phasic pupil dilation in response to target sounds and facilitated detection. By contrast, the present study aimed to *increase* tonic arousal by showing highly arousing video and music clips. The absence of an effect of tonic arousal modulation on phasic physiological responses and behavioral performance may be explained by methodological aspects. Indeed, isometric handgrip is a more direct and physiologically intense method for eliciting NE release, while music and video may not trigger the same physiological cascade or intensity of LC-NE engagement. Moreover, a systematic review highlights that music-induced changes in autonomic arousal measures (HR, SCL, PD) and cognitive performance are highly variable across individuals, influenced by factors such as personal preference and the role of music in daily life (Chee et al., 2024). In the present study, this interindividual variability likely added noise to our measures across participants and diluted potential small effects.

Additionally, the timing of the tonic arousal effect may have limited our ability to detect its influence on phasic responses and behavior. As discussed above, the effect of tonic arousal manipulation was evidenced only for the first part of the oddball task; similar to Mather et al. (2020). Since our analysis of phasic trials considered the entire task duration, a transient early effect on phasic measures might have been diluted. An additional analysis of trials from the time-windows showing heightened tonic arousal (first half of trials in each block, ∼87 s) confirmed that there was indeed no induction effect.

Finally, theoretical accounts may explain why no tonic–phasic relationship emerged here. Our hypotheses regarding the influence of tonic arousal on phasic responses were originally based on the proposed inverted-U-relationship between arousal and performance (Aston-Jones & Cohen, 2005; Yerkes & Dodson, 1908), which predicts that heightened tonic activity can impact phasic responses and impair performance. Contrary to this expectation, our results showed no evidence that increases in tonic arousal affected phasic responses and performance. One likely explanation is task difficulty: according to the unification model of attention with the Adaptive Gain Theory and the GANE model (R. Huang & Clewett, 2024), the classic inverted-U relationship mainly applies to cognitively demanding (i.e. difficult) tasks. In easier tasks, like the one used here, rising tonic arousal may enhance processing without impacting phasic responses, consistent with a monotonic arousal-performance relationship. It remains possible that at higher arousal levels or with more challenging tasks, tonic-phasic interactions would emerge, which should be systematically examined with the present approach in the future.

### Developmental differences

#### Behavioral evidence of immature attentional control in 6–8 year-old children

Several studies have demonstrated developmental improvements in attentional task performance, with adults showing higher accuracy and faster response times compared to children, likely reflecting the maturation of attentional control processes (e.g., Wetzel et al., 2016, 2021). Our results align with this general pattern: children made more false alarms and responded more slowly than adults. Notably, only a few studies have used an active three-stimulus auditory oddball paradigm in children, and they report similar developmental trends in both accuracy and reaction time (Cycowicz et al., 1996; Määttä, Pääkkönen, et al., 2005; Mingils et al., 2023), supporting the robustness of these effects.

Additionally, we found a developmental effect on reaction time variability, considered a robust measure of sustained attention (Antonini et al., 2013). Increased reaction time variability is typically observed in both children and adults with ADHD (Kofler et al., 2013; Tamm et al., 2012). In our typically developing sample, children exhibited greater reaction time variability than adults, consistent with prior findings in the general population showing reduced abilities in sustained attention in children (Dykiert et al., 2012; Hoyer et al., 2021; Williams et al., 2005).

Together, these behavioral results strengthen the evidence for ongoing maturation of attentional functions across childhood, reflected in both mean performance and intra-individual variability.

#### Children show higher tonic arousal and greater sensitivity to highly arousing videos

Besides behavioral differences, we observed developmental effects on two physiological markers of tonic arousal. First, children showed larger pupil diameters than adults, consistent with previous findings suggesting stronger sympathetic nervous system (SNS) activity and/or weaker parasympathetic nervous system (PNS) influence in younger populations (Bufo et al., 2022; Tekin et al., 2018). However, pupil diameter development appears non-linear: some studies report larger pupil diameters in children; whereas others describe a progressive increase during childhood and adolescence, followed by a decline in adulthood (MacLachlan & Howland, 2002; Silbert et al., 2013; Winston et al., 2020). This trajectory likely reflects both autonomic nervous system maturation and anatomical changes such as in the iris volume (Jouzdani et al., 2013).

Second, children displayed higher skin conductance level than adults. This effect is also likely to be multifactorial. On one hand, skin conductance is influenced by physiological properties of the skin, such as thickness and sweat gland density, which vary with age and can affect the skin’s electrical characteristics (Boucsein, 1992). On the other hand, it reflects SNS activity, which may exhibit greater reactivity in children (Bufo et al., 2022).

Interestingly, children showed stronger skin conductance levels to the highly arousing videos than adults, during the second half of the induction period. This specific effect of highly arousing videos cannot be related to physiological properties of the skin, but rather suggest higher reactivity of the SNS in children. This heightened reactivity may reflect greater sensitivity to the video content, which consisted of short, humorous animated clips designed for a young audience, onto which excerpts of high-arousing music were superimposed. Subjective ratings support this interpretation: children perceived the highly arousing videos as more stimulating than the music alone, while adults rated both types of stimuli as similarly arousing. In contrast, adults may require more intense or emotionally charged content, such as unpleasant or stressful stimuli, to elicit comparable levels of autonomic arousal (Lang et al., 1993). However, such stimuli may not be appropriate for use with children. These considerations highlight a methodological challenge in developmental research: identifying stimuli that are both engaging and ethically appropriate for children while also comparably arousing across age groups.

Importantly, we observe a progressive decrease in SCL during the task in adults, while SCL stays relatively stable in children. In adults, SCL often shows a gradual decline (habituation or adaptation) during prolonged, monotonous, or non-arousing task blocks, reflecting reduced sympathetic arousal as the environment or task becomes less novel (Barry & Sokolov, 1993). This habituation response is also shown in fMRI studies with decreased amygdala activation during repeated presentation of emotional stimuli (Breiter et al., 1996). The absence of decline in children’s SCL throughout the task block (approx. 3 min) might index that children were more motivated or aroused by the task than adults (Malmo, 1965), irrespectively of the induction condition. Also, in children, higher tonic arousal may help maintain task performance by mitigating challenges in sustaining attention.

#### Children display heightened phasic responses to task-irrelevant sounds

A major interest of the present study was to examine phasic physiological responses (SCR, PDR) to task-relevant and irrelevant sounds during an active three-stimulus oddball task in children. We found strong developmental effects on those phasic responses.

Overall, SCRs were larger in children than in adults. Both groups showed stronger responses to targets than to novels, and to novels than to standard stimuli, but this effect was amplified in children. Indeed, children presented larger SCR in response to both relevant (targets) and irrelevant (novels) rare salient sounds compared to adults. Our findings show that in children, rare salient auditory events trigger a stronger enhancement of sympathetic activation, as indexed by SCR. These larger responses in phasic arousal to unexpected events could be related to increased attentional capture in children, compared to adults.

Unlike skin conductance, which is under the unique control of the SNS, pupil dilation is innervated by both SNS and PNS, with the early peak reflecting parasympathetic inhibition and the late peak sympathetic activation (Widmann et al., 2018). Different patterns of activation were found for each peak. The early peak of the PDR was larger in children than in adults, but only for task-irrelevant sounds. In adults, the early peak was largest for targets, smaller for novels, and smallest for standards, whereas in children the early response to novels was as large as the response to targets. Thus, children present strong early reactivity to both types of rare stimuli, irrespective of task relevance. The late peak of the PDR was larger for targets than for novels, and larger for novels than for standards, with no difference between groups. In children, the absence of differentiation between targets and novels in the early peak suggests that the parasympathetic system first responds strongly to all non-standard stimuli, with task-based differentiation emerging only at the late, sympathetic-driven response. In adults, by contrast, task-relevant information is already filtered at the early peak. This appears consistent with SCR results, showing differential sympathetic-driven responses to novel and target in both children and adults. These findings indicate that attentional control is still developing in children, with top-down selection of relevant versus irrelevant cues occurring later in time than in adults.

Similar developmental differences in the processing of novel sounds have been reported in EEG studies using three-stimulus oddball paradigms, with children showing P3 responses to novel stimuli that differ from adults particularly in scalp distribution (Cycowicz et al., 1996; Määttä, Saavalainen, et al., 2005; Wetzel & Schröger, 2007, 2014). Such findings are in line with developmental accounts suggesting that children show greater reactivity to novel or unexpected stimuli, potentially due to less efficient attentional control and immature arousal systems (Pozuelos et al., 2014; Wetzel & Schröger, 2014).

The present physiological results provide new insight into children’s attentional functioning. Children show heightened responsiveness to both task-relevant and irrelevant stimuli compared to adults, which may contribute to their greater distractibility. This increased distractibility has been linked in behavioral studies to both increased sensitivity to irrelevant input and an ongoing maturation of voluntary attentional control (Hoyer et al., 2021).

While adults outperform young children in most behavioral tasks, this increased sensitivity to bottom-up inputs can lead in specific cases to a developmental reversal in attention, where children outperform adults in paying attention or remembering salient stimuli in change-detection or incidental learning tasks (Plebanek & Sloutsky, 2017; Tandoc et al., 2024). This broader attentional distribution may thus support exploratory behavior, critical for learning (Blanco & Sloutsky, 2020; Plebanek & Sloutsky, 2017). By showing that children exhibit larger phasic arousal responses to task-irrelevant stimuli, the present study provides physiological evidence that this broader attentional focus may promote exploration, in line with Aston-Jones’ framework linking arousal to the exploration–exploitation trade-off (Aston-Jones & Cohen, 2005).

## Conclusion

Taken together, our findings contribute to the understanding of how children and adults respond to rare, salient auditory stimuli. Children exhibited heightened phasic responses, particularly to task-irrelevant novel sounds, reflecting a less mature regulation of arousal-attention interactions. However, despite successful external modulation of tonic arousal, phasic responses remained unaffected, suggesting that tonic arousal exerts limited influence on stimulus-driven responses in a simple attention task. Future studies could explore more challenging tasks and stronger arousal manipulations to better reveal potential tonic–phasic interactions. Finally, our study demonstrates the potential of non-invasive techniques, such as skin conductance and pupil dilation, to investigate the interplay between arousal and attention in children, offering promising tools for future developmental research.

