## Supplementary Figures for "Tuned to explore: Increased phasic responses to auditory targets and novelty in children regardless of induced tonic arousal"

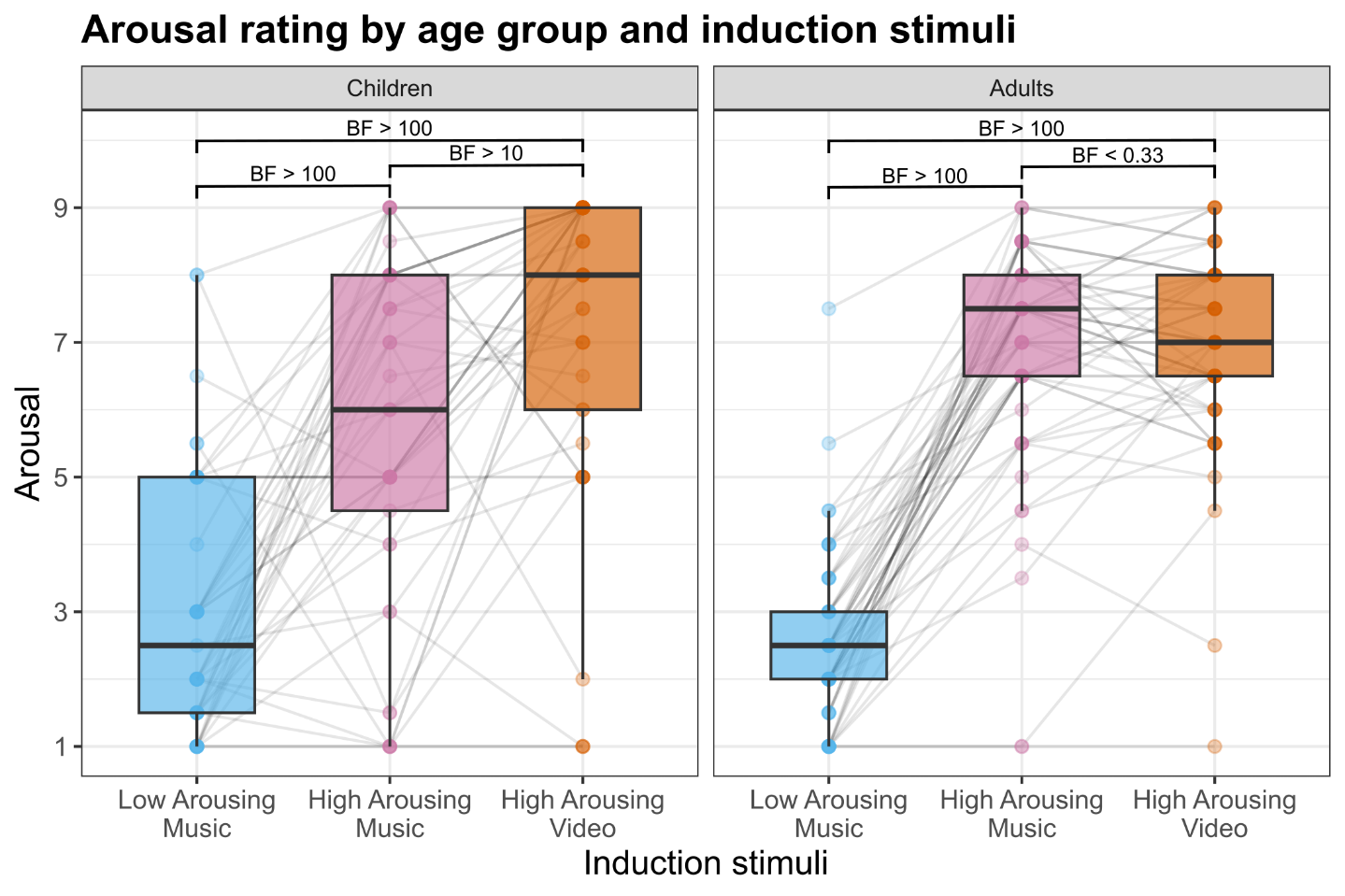

**Sup. Fig. 1.** Arousal ratings by age and induction stimuli. Within each boxplot, the horizontal line represents the group median, the lower and upper hinges correspond to the first and third quartiles. The upper whisker extends from the hinge to the largest value no further than 1.5 * IQR from the hinge (IQR = inter-quartile range, or distance between the first and third quartiles). The lower whisker extends from the hinge to the smallest value at most 1.5 * IQR of the hinge. Superimposed to each boxplot, the dots represent individual means. Light grey lines connect the dots belonging to the same participant across conditions. BF: Bayes Factor.

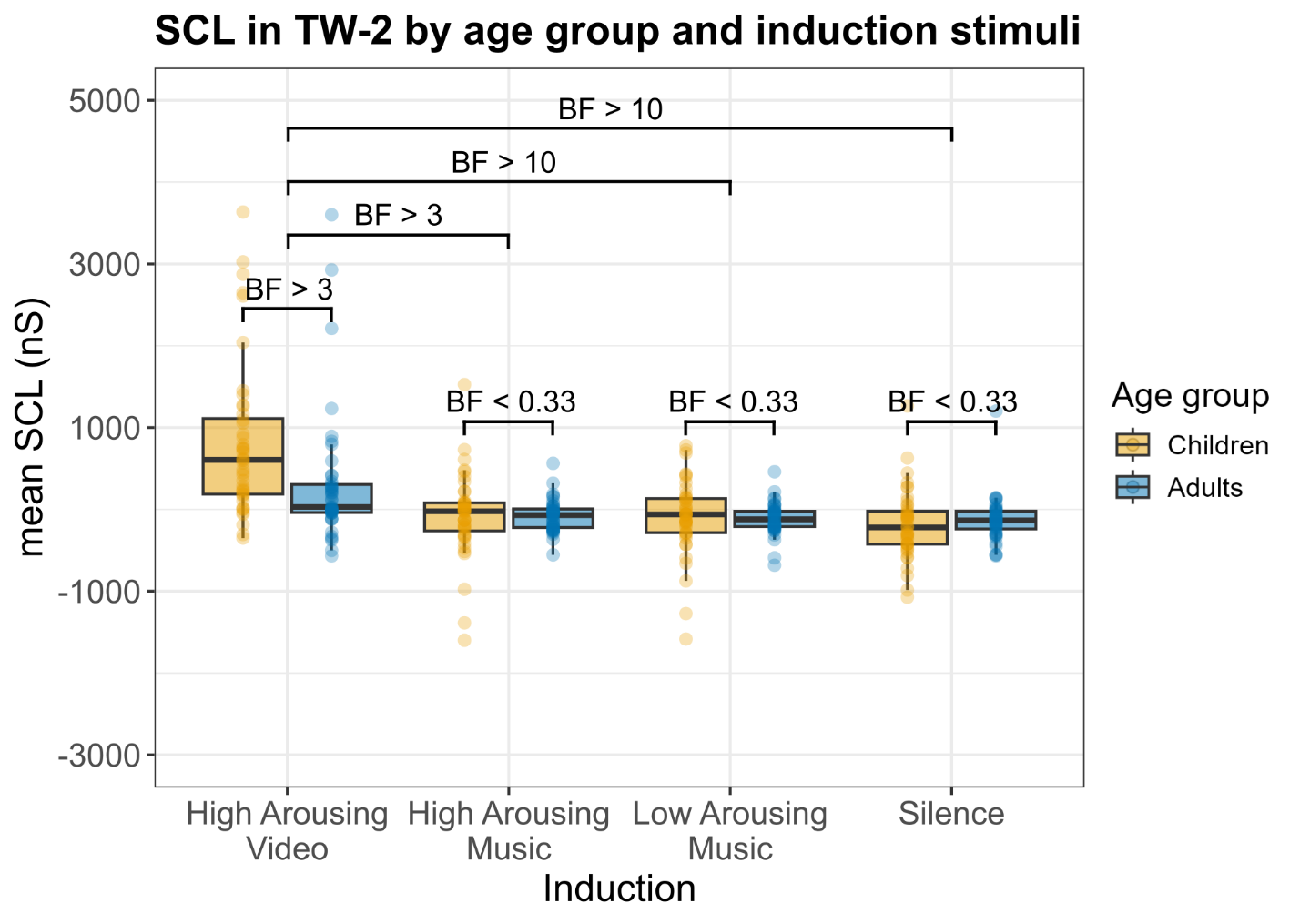

**Sup. Fig. 2**. Mean Skin conductance level (SCL) in TW-2 as a function of age group and induction. Within each boxplot, the horizontal line represents the group median, the lower and upper hinges correspond to the first and third quartiles. The upper whisker extends from the hinge to the largest value no further than 1.5 * IQR from the hinge (IQR = inter-quartile range, or distance between the first and third quartiles). The lower whisker extends from the hinge to the smallest value at most 1.5 * IQR of the hinge. Superimposed to each boxplot, the dots represent individual means. BF: Bayes Factors.

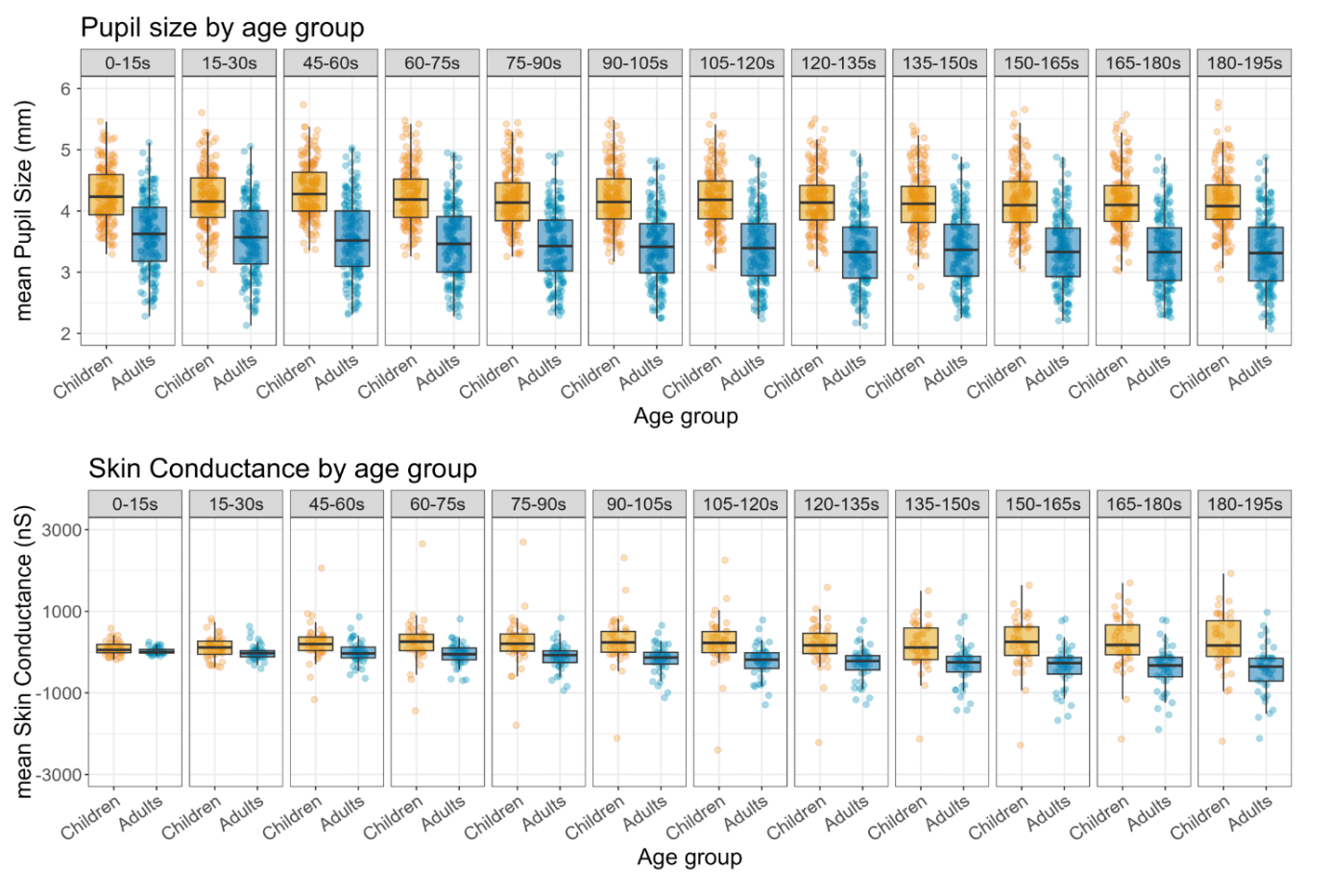

**Sup. Fig. 3**. Pupil size by age group for each 15-second TWs corresponding to the induction period (0-30s) and the following oddball task (45-195s). Within each boxplot, the horizontal line represents the group median, the lower and upper hinges correspond to the first and third quartiles. The upper whisker extends from the hinge to the largest value no further than 1.5 * IQR from the hinge (IQR = inter-quartile range, or distance between the first and third quartiles). The lower whisker extends from the hinge to the smallest value at most 1.5 * IQR of the hinge. Superimposed to each boxplot, the dots represent individual means.

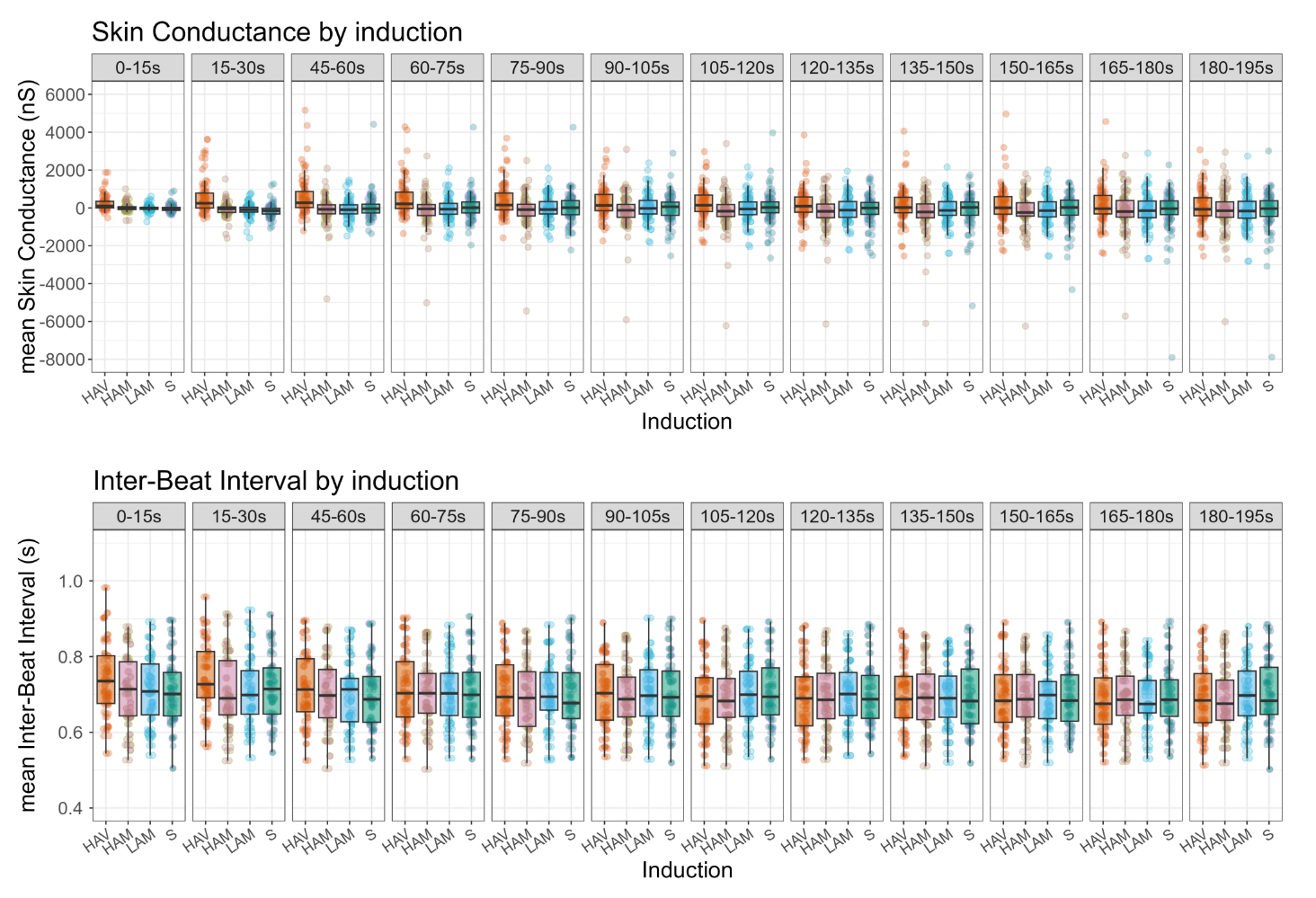

**Sup. Fig. 4**. Inter-Beat Interval by induction condition for each 15-second TWs corresponding to the induction period (0-30s) and the following oddball task (45-195s). Within each boxplot, the horizontal line represents the group median, the lower and upper hinges correspond to the first and third quartiles. The upper whisker extends from the hinge to the largest value no further than 1.5 * IQR from the hinge (IQR = inter-quartile range, or distance between the first and third quartiles). The lower whisker extends from the hinge to the smallest value at most 1.5 * IQR of the hinge. Superimposed to each boxplot, the dots represent individual means.
