## Supplementary Tables for "Tuned to explore: Increased phasic responses to auditory targets and novelty in children regardless of induced tonic arousal"

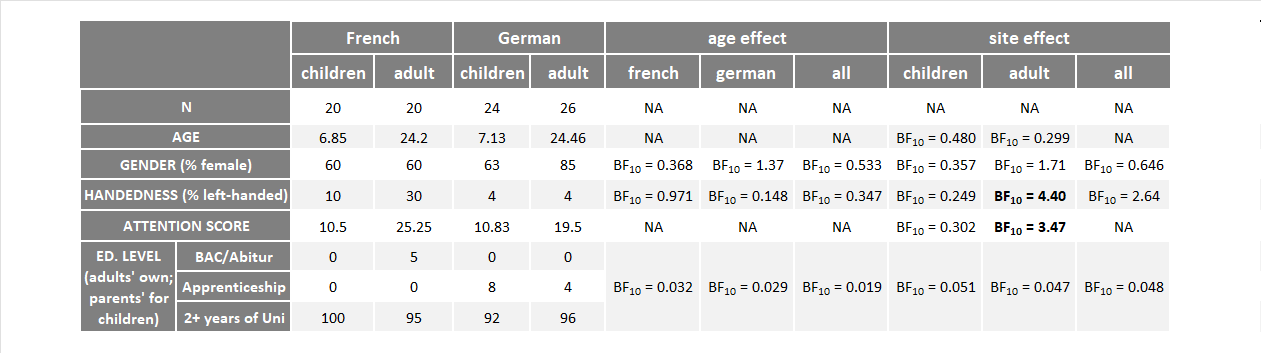

**Sup. Table 1.** Demographic characteristics of participants. Age effect refers to differences between children and adults within each site and across all participants. Site effect refers to differences between the French and German recording sites within each age group and across all participants.

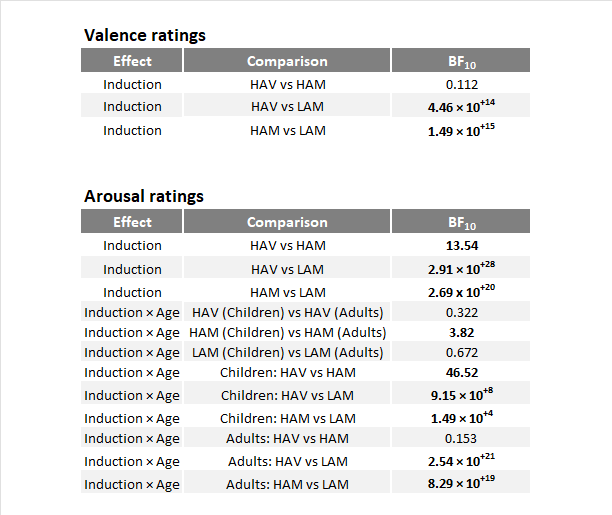

**Sup. Table 2.** Results of Bayesian post-hoc t-tests for valence (top) and arousal (bottom) ratings of the induction stimuli. HAV: High Arousing Video; HAM: High Arousing Music; LAM: Low Arousing Music. BF: Bayes Factor.

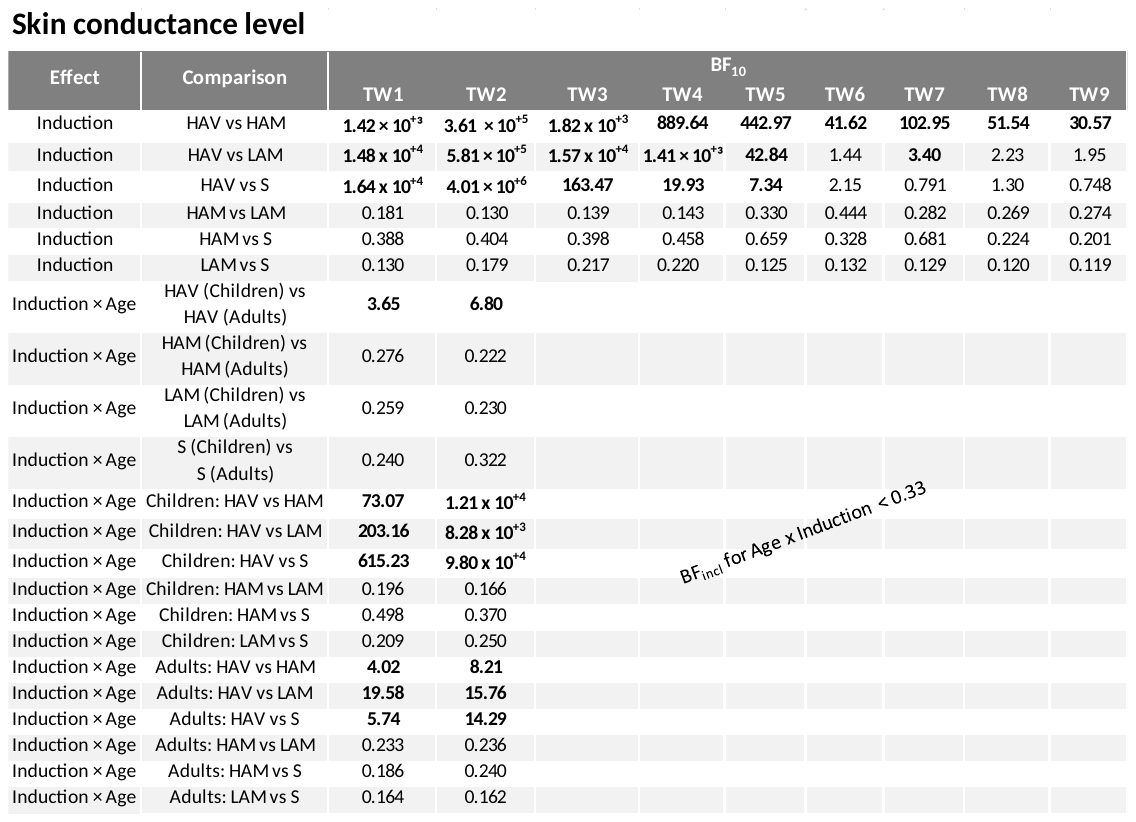

**Sup. Table 3.** Results of Bayesian post-hoc t-tests for skin conductance level. HAV: High Arousing Video; HAM: High Arousing Music; LAM: Low Arousing Music; S: Silence. BF: Bayes Factor.

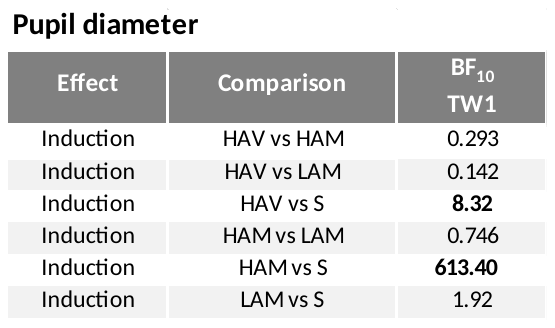

**Sup. Table 4.** Results of Bayesian post-hoc t-tests for pupil diameter. HAV: High Arousing Video; HAM: High Arousing Music; LAM: Low Arousing Music; S: Silence. BF: Bayes Factor.

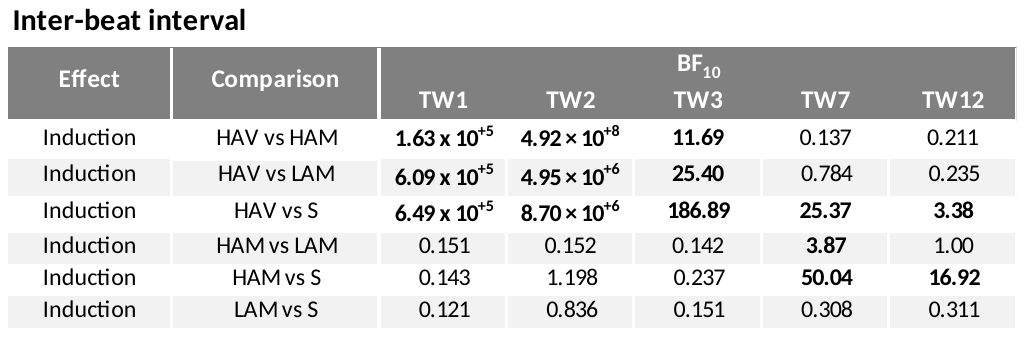

**Sup. Table 5.** Results of Bayesian post-hoc t-tests for inter-beat interval. HAV: High Arousing Video; HAM: High Arousing Music; LAM: Low Arousing Music; S: Silence. BF: Bayes Factor.

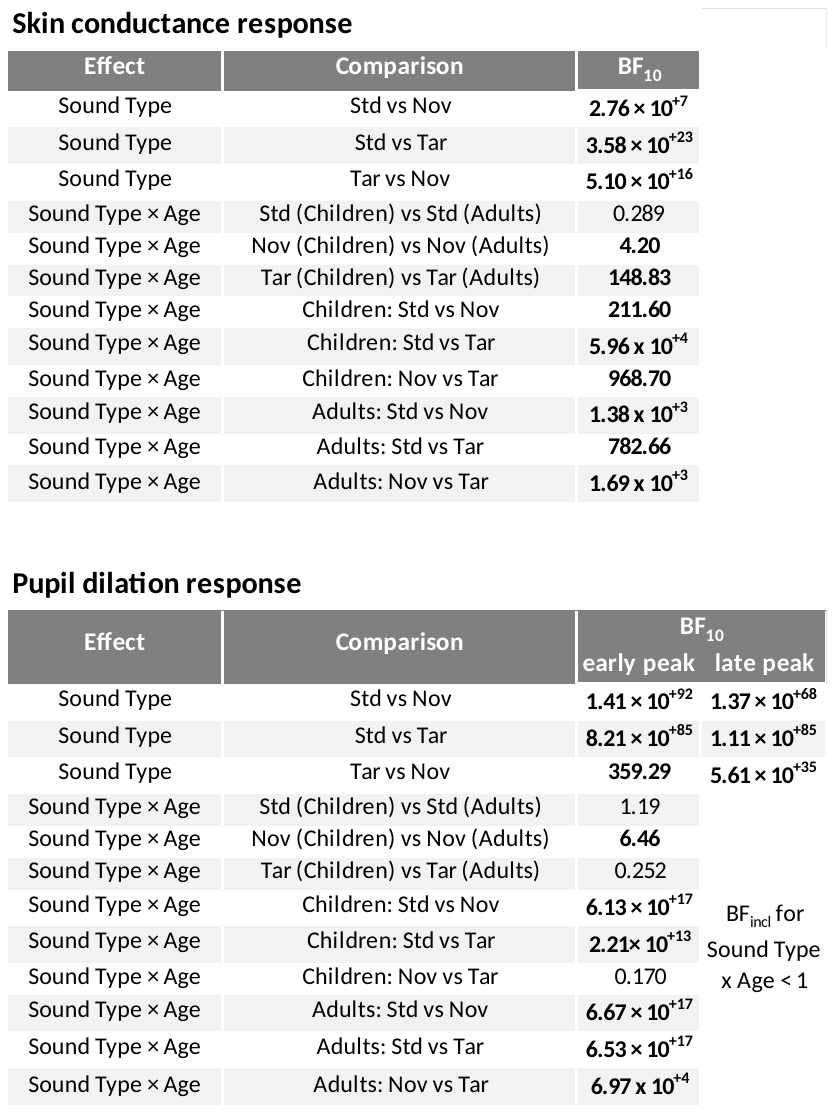

**Sup. Table 6.** Results of Bayesian post-hoc t-tests for phasic responses: skin conductance response (**top**) and pupil dilation response (**bottom**). Std: standard sound; Nov: novel sound; Tar: target sound. BF: Bayes Factor.

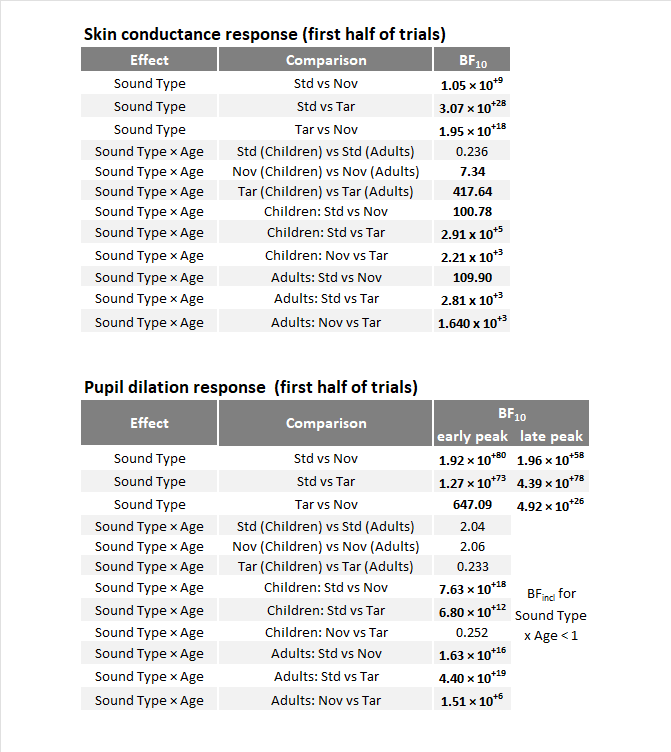

**Sup. Table 7.** Results of Bayesian post-hoc t-tests for phasic responses in the first half of the trials: skin conductance response (**top**) and pupil dilation response (**bottom**). Std: standard sound; Nov: novel sound; Tar: target sound. BF: Bayes Factor.
